# Cocaine sensitization and accumbens shell plasticity depend on biological sex and gonadal hormones in C57BL/6J mice

**DOI:** 10.64898/2025.12.11.693721

**Authors:** Andrew D. Chapp, Chinonso A. Nwakama, Chau-Mi H. Phan, Pramit P. Jagtap, Erin B. Lind, Andréa R. Collins, Yanaira Alonso-Caraballo, Mark J. Thomas, Paul G. Mermelstein

**Affiliations:** Department of Neuroscience, University of Minnesota, Minneapolis, MN 55455; Medical Discovery Team on Addiction, University of Minnesota, MN 55445; Medical Scientist Training Program, Icahn School of Medicine at Mount Sinai, New York, NY 10029; Nash Family Department of Neuroscience, Friedman Brain Institute, Icahn School of Medicine at Mount Sinai, New York, NY 10029; Department of Psychiatry, University of California, San Francisco Fresno, Fresno, CA 93701; Center for Neural Circuits in Addiction, University of Minnesota, Minneapolis, MN 55455

## Abstract

Biological sex as a defining variable in drug sensitivity remains poorly understood. Here, we combine behavioral and electrophysiological analyses to examine the influence of sex and gonadal hormones on cocaine-induced psychomotor sensitization and nucleus accumbens shell (NAcSh) plasticity in the prominent C57BL/6J mouse strain. Males exhibited greater cocaine-evoked locomotor activity than females; castration attenuated responses, whereas ovariectomy enhanced them. This behavioral phenotype is opposite to what occurs in rats. A 10–14 day abstinence period abolished the sex difference in intact animals, and gonadectomy reduced cocaine-induced behavioral plasticity. Recordings from 309 medium spiny neurons revealed sex-dependent NAcSh plasticity. In males, cocaine decreased neuronal excitability, while in females it induced estrous cycle–dependent plasticity characterized by reduced excitability during diestrus relative to estrus. These effects were driven by cocaine-induced modulation of voltage-gated sodium channels. Cocaine potentiated glutamatergic strength in males but elicited estrous cycle–dependent depotentiation in females. These adaptations in excitability and glutamatergic strength were abolished by gonadectomy, and paralleled diminished behavioral plasticity during abstinence. These data illustrate that biological sex and hormonal milieu critically shape cocaine-induced plasticity, offering a more nuanced framework than the traditional notion of heightened female sensitivity to drugs of abuse.

## INTRODUCTION

Classically, rodent models of addiction exploring sex differences in drug sensitivity frequently report greater drug effects in females compared to males. For cocaine, females generally have increased locomotion, escalation, and relapse compared to males^1, 2, 3, 4, 5, 6, 7, 8^. The estrous cycle and ovarian hormones have been implicated in these enhanced behavioral responses to cocaine^7, 9, 10, 11, 12, 13^. Estradiol, produced by the ovaries in females is the primary sex hormone thought to be responsible for this enhancement^10, 11, 12, 13, 14, 15, 16^. However, unlike much of the consistency in rodent models (i.e. primarily rats), human studies have reported mixed results^17, 18, 19^. Consistent with female rats being more sensitive than males, the menstrual cycle has been shown to alter the subjective feelings of cocaine exposure^19, 20^ as well as cravings for cocaine^19, 20^. However, a recent systematic review has revealed there is little evidence for sex differences in stimulant use in humans^17^. This suggests that there are potential caveats to sex-specific sensitivities in psychostimulants, such as cocaine, and finding these exceptions may be crucial to advancing our understanding of cocaine use disorder (CUD) among males and females.

We have previously shown that the genetic background of mice has a profound effect on the sex differences in cocaine psychomotor sensitization^21^. Specifically, in the extensively utilized C57BL/6J (C57) strain, male animals had increased cocaine psychomotor sensitization compared to females^21^. Given the widespread use of C57s in cocaine research and the lack of electrophysiology data in females during abstinence in this strain, we prioritized understanding how biological sex and gonadal hormones affect cocaine behavior and nucleus accumbens shell (NAcSh) medium spiny neuron (MSN) electrophysiology during abstinence.

As addiction is believed to be driven by drug-induced neuroplastic changes in reward areas like the NAcSh, understanding cocaine’s role in shaping abstinence neuroplasticity in this brain region is crucial for predicting vulnerabilities and treatments for CUD which may be sex-specific. Importantly, cocaine plasticity intersects with estrous cycle plasticity in females, which requires estrous cycle tracking to determine potential cycle-specific effects of cocaine^7, 22, 23^. Here, using cocaine behavioral psychomotor sensitization paired with *ex vivo* electrophysiology, we uncover sex and gonadal hormone differences in both cocaine sensitization and NAcSh cocaine abstinence plasticity-findings which fundamentally challenge our current understanding of cocaine neuropharmacology.

## MATERIALS AND METHODS

### Animals

Animal procedures were performed at the University of Minnesota in facilities accredited by the Association for Assessment and Accreditation of Laboratory Animal Care (AAALAC) and in accordance with protocols approved by the University of Minnesota Institutional Animal Care and Use Committee (IACUC), as well as the principles outlined in the National Institutes of Health *Guide for the Care and Use of Laboratory Animals*. Male and female mice C57BL/6J (#000664), including castrated (CAST) males and ovariectomized (OVX) females were obtained from Jackson Labs (Bar Harbor, ME). Castration and ovariectomy surgeries were performed when mice were 3 weeks of age or older and were performed at Jackson Labs (Bar Harbor, ME), or a portion of the ovariectomy surgeries were performed at the University of Minnesota. Mice aged 8-12 weeks were used in all experiments, were group housed and kept on a 10:14 dark:light cycle with food and water *ad libitum*. One hundred thirty-five (135) animals were used for behavioral sensitization: male cocaine (34), male saline (6), female cocaine (39), female saline (8), OVX cocaine (17), OVX saline (6), CAST cocaine (18), and CAST saline (7). A subset of the animals underwent a challenge cocaine injection: Female cocaine (15), male cocaine (18), CAST cocaine (9), and OVX cocaine (9). Electrophysiology for male cocaine (8), male saline (7), female cocaine (15), and female saline (15) animals were obtained from animals that underwent cocaine behavioral sensitization reported in Chapp et al^21^. Electrophysiology in GDX animals were obtained from: OVX cocaine (6), OVX saline (6), CAST cocaine (6), and CAST saline (6).

### Ovariectomy Surgery

Bilateral ovariectomies were performed in aseptic conditions under isoflurane anesthesia (3%-5% pre-operative induction, 1%-2% intra-operative maintenance) similar to previously described^24^. On each side, a dermal incision was made to the flank region just below the rib cage. Forceps were used to separate the underlying connective tissue and to penetrate through the muscle layer into the peritoneal cavity. The ovarian fat pad was then located, gently externalized, and forceps were used to locate the distal uterine horn, oviduct, and ovary. The distal end of the uterine horn was ligated with suture and the ovary subsequently removed using surgical scissors. The remaining tissue was then returned to the abdominal cavity, the muscle wall resealed with suture, and the dermal incision closed with suture and wound clips (7mm). Carprofen (10 mg/kg, s.c.) was given for pre- and post-operative analgesia, and wound clips were removed after 10 days. Successful ovariectomy was confirmed by vaginal lavage at the end of study.

### Psychomotor sensitization

All experiments were conducted between 12:00 and 17:00 hrs, with houselights on at 06:00 and off at 20:00 hrs. Animals were handled and habituated to locomotor chambers as well as subcutaneous (S.C.) injections two days prior to behavioral testing. On test days, animals were given either an S.C. injection of cocaine (15 mg/kg) daily for five days^25^, or an equivalent volume of 0.9% saline and placed promptly into the behavioral testing chamber (18” x 9”, with 8.5” tall walls) with light levels of 250 ± 10 lux. Videos were recorded for 80 minutes using ANY-maze tracking software (Wood Dale, IL) and the total distance traveled was analyzed offline.

### Chemicals

All chemicals were obtained from Sigma-Aldrich (St Louis, MO, USA), except cocaine hydrochloride and isoflurane (Boynton Pharmacy, University of Minnesota, MN, USA) and carprofen (Research Animal Resources, University of Minnesota, MN, USA).

### Whole-cell recordings

Mice (8-12 weeks old) in abstinence (10-14 days after the last behavioral day) were used for electrophysiology recordings. Animals were sacrificed between 09:00 and 17:00. For females, prior to being anesthetized, estrous cycle was determined by vaginal cytology as previously described ^26, 27^. Mice were anesthetized with isoflurane (3% in O_2_) and decapitated. The brain was rapidly removed and chilled in ice cold cutting solution, containing (in mM): 228 sucrose, 2.5 KCl, 7 MgSO_4_, 1.0 NaH_2_PO_4_, 26 NaHCO_3_, 0.5 CaCl_2_, 11 d-glucose, pH 7.3-7.4, continuously gassed with 95:5 O_2_:CO_2_ to maintain pH and pO_2_. A brain block was cut including the NAcSh region and affixed to a vibrating microtome (Leica VT 1000S; Leica, Nussloch, Germany). Sagittal sections of 240 µm thickness were cut, and the slices transferred to a holding container of artificial cerebral spinal fluid (ACSF) maintained at 30 °C, continuously gassed with 95:5 O_2_:CO_2_, containing (in mM): 119 NaCl, 2.5 KCl, 1.3 MgSO_4_, 1.0 NaH_2_PO_4_, 26.2 NaHCO_3_, 2.5 CaCl_2_, 11 d-glucose, and 1.0 ascorbic acid (osmolality: 295–302 mOsmol L−1; pH 7.3-7.4)28, 29 and allowed to recover for 1 hr. Following recovery, slices were transferred to a glass-bottomed recording chamber and viewed through an upright microscope (Olympus) equipped with DIC optics, and an IR-sensitive video camera (DAGE-MTI).

Slices transferred to the glass-bottomed recording chamber were continuously perfused with ACSF, gassed with 95:5 O_2_:CO_2_, maintained at room temperature and circulated at a flow of 2 mL min^-1^. Patch electrodes were pulled (Flaming/Brown P-97, Sutter Instrument, Novato, CA) from borosilicate glass capillaries with a tip resistance of 5–10 MΩ. Electrodes were filled with a solution containing (in mM) 135 K-gluconate, 10 HEPES, 0.1 EGTA, 1.0 MgCl_2_, 1.0 NaCl, 2.0 Na_2_ATP, and 0.5 Na_2_GTP (osmolality: 280–285 mOsmol L^−1^; pH 7.3)^30^. MSNs were identified under IR-DIC based on morphology and their hyperpolarizing membrane potential (-70 to -80 mV) and were voltage clamped at -80 mV using a Multiclamp 700B amplifier (Molecular Devices), currents filtered at 2 kHz and digitized at 10 kHz. Holding potentials were not corrected for the liquid junction potential. Once a GΩ seal was obtained, slight suction was applied to break into whole-cell configuration and the cell was allowed to stabilize which was determined by monitoring capacitance, membrane resistance, access resistance and resting membrane potential (V_m_)^30, 31, 32^. Cells that met the following criteria were included in the analysis: action potential amplitude ≥50 mV from threshold to peak, resting *V*_m_ negative to −64 mV, and <20% change in series resistance during the recording. Passive membrane properties, capacitance and membrane resistance were measured from the membrane test in pClamp (Molecular Devices). Resting neuronal membrane potential reported in Table 1 was recorded immediately after breaking into whole-cell mode^22, 27, 30, 33, 34^. To measure NAcSh MSN neuronal excitability, V_m_ was adjusted to -80 mV by continuous negative current injection, and a series of square-wave current injections was delivered in steps of +20 pA, each for a duration of 800 ms. To determine the action potential voltage threshold (Vt), and rheobase, ramp current injections (0.437 pA/ms, 800 ms) were made from a holding potential of -80 mV.

**Table 1.** Passive NAcSh MSN membrane properties.

|  | **Capacitance (pF)** | **V<sub>m</sub> (mV)** | **R<sub>m</sub> (MΩ)** |
| --- | --- | --- | --- |
| **Female estrus saline (n=18)** | 87.6 ± 2.8 | -77.8 ± 0.9 | 95.9 ± 4.6 |
| **Female estrus cocaine (n=16)** | 83.6 ± 2.8 | -77.7 ± 0.7 | 103.6 ± 3.8 |
| P value | 0.608 | 0.999 | 0.999 |
| **Female diestrus saline (n=18)** | 84.3 ± 2.5 | -76.0 ± 0.8 | 95.7 ± 4.5 |
| **Female diestrus cocaine (n=18)** | 84.3 ± 3.0 | -77.4 ± 0.6 | 100.2 ± 3.3 |
| P value | 0.999 | 0.999 | 0.999 |
| **OVX Female saline (n=13)** | 86.5 ± 3.3 | -74.1 ± 0.7 | 106.0 ± 6.1 |
| **OVX Female cocaine (n=13)** | 86.7 ± 2.1 | -74.2 ± 0.8 | 104.3 ± 4.7 |
| P value | 0.999 | 0.999 | 0.999 |
| **Male saline (n=25)** | 74.0 ± 3.6 | **-73.2 ± 1.0** | **124 ± 10.2** |
| **Male cocaine (n=15)** | 80.2 ± 3.5 | **-76.9 ± 0.7** | **100.7 ± 4.7** |
| P value | 0.84 | **0.02** | **0.02** |
| **CAST male saline (n=29)** | 77.9 ± 2.4 | -75.3 ± 0.8 | 102.8 ± 3.9 |
| **CAST male cocaine (n=20)** | 76.0 ± 3.1 | -74.5 ± 1.0 | 94.5 ± 3.5 |
| P value | 0.99 | 0.99 | 0.99 |

Abbreviations: V<sub>m</sub>= Resting membrane potential, R<sub>m</sub>= membrane resistance, OVX=ovariectomy CAST= castrated.

For miniature excitatory postsynaptic current recordings (mEPSC)^33^, slices transferred to the glass-bottomed recording chamber were continuously perfused with ACSF containing lidocaine (0.7 mM) to block voltage-gated sodium channels and picrotoxin (100 μM) to block GABAR, and was continuously gassed with 95:5 O_2_:CO_2_, maintained at room temperature and circulated at a flow of 2 mL min^-1^. Patch electrodes were pulled from borosilicate glass capillaries with a tip resistance of 5– 10 MΩ and whole-cell recordings were made. Electrodes were filled with a cesium methanesulfonate (CsMeSO_4_) solution containing (in mM): 120 CsMeSO4, 15 CsCl, 10 TEA-Cl, 10 HEPES, 0.4 EGTA, 8.0 NaCl, 2.0 Na2ATP, and 0.3 Na2GTP (osmolality: 280–285 mOsmol L−1; pH 7.3). MSNs were identified under IR-DIC based on morphology and their hyperpolarizing membrane potential (-70 to -80 mV) and were voltage clamped at -80 mV using A Multiclamp 700B amplifier (Molecular Devices), currents filtered at 2 kHz and digitized at 10 kHz. Holding potentials were not corrected for the liquid junction potential. mEPSCs were recorded for 2 minutes and analyzed offline using Mini Analysis software (synaptosoft) with an amplitude threshold set at three times the noise level.

### Statistical analysis

Data values were reported as mean ± SEM. All statistical analyses were performed with a commercially available statistical package (GraphPad Prism, version 9.4.1). Probabilities less than 5% were deemed significant *a priori*. Depending on the experiments, group means were compared using an unpaired Student’s *t*-test, a one-way ANOVA, or a one- or two-way ANOVA with repeated measures or mixed effects. Where differences were found, Fisher’s LSD post hoc tests were used for multiple pair-wise comparisons.

## RESULTS

### Male mice have increased cocaine-induced locomotion compared to female mice

We previously reported sex-dependent differences in cocaine (15 mg/kg) psychomotor sensitization in C57BL/6J mice (males>females)^21^. In cocaine-treated animals utilizing this identical sensitization paradigm (Fig 1A), we replicated our previous findings where male mice displayed an overall increase in cocaine locomotion compared to female mice (Fig 1B; two-way ANOVA-Mixed, F_(1,71)_=6.284, p<0.05). Males sensitized to cocaine (days 3-7, one-way ANOVA-RM, F_(2.73,90.05)_=165.1, p<0.0001) as did females (days 3-7, one-way ANOVA-RM, F_(3.08,116.9)_=176.7, p<0.0001). The addition of a cocaine challenge after 10-14 days of abstinence abolished the sex-dependent differences in cocaine psychomotor sensitization (Fig 1B,C; two-way ANOVA-Mixed, F_(1,31)_=0.15, p=0.698). This was due to a slight loss in the cocaine psychomotor effects in males and an increase in the psychomotor effects in females, suggesting a potential difference in cocaine sensitivity between the sexes following abstinence.

**Fig 1.**
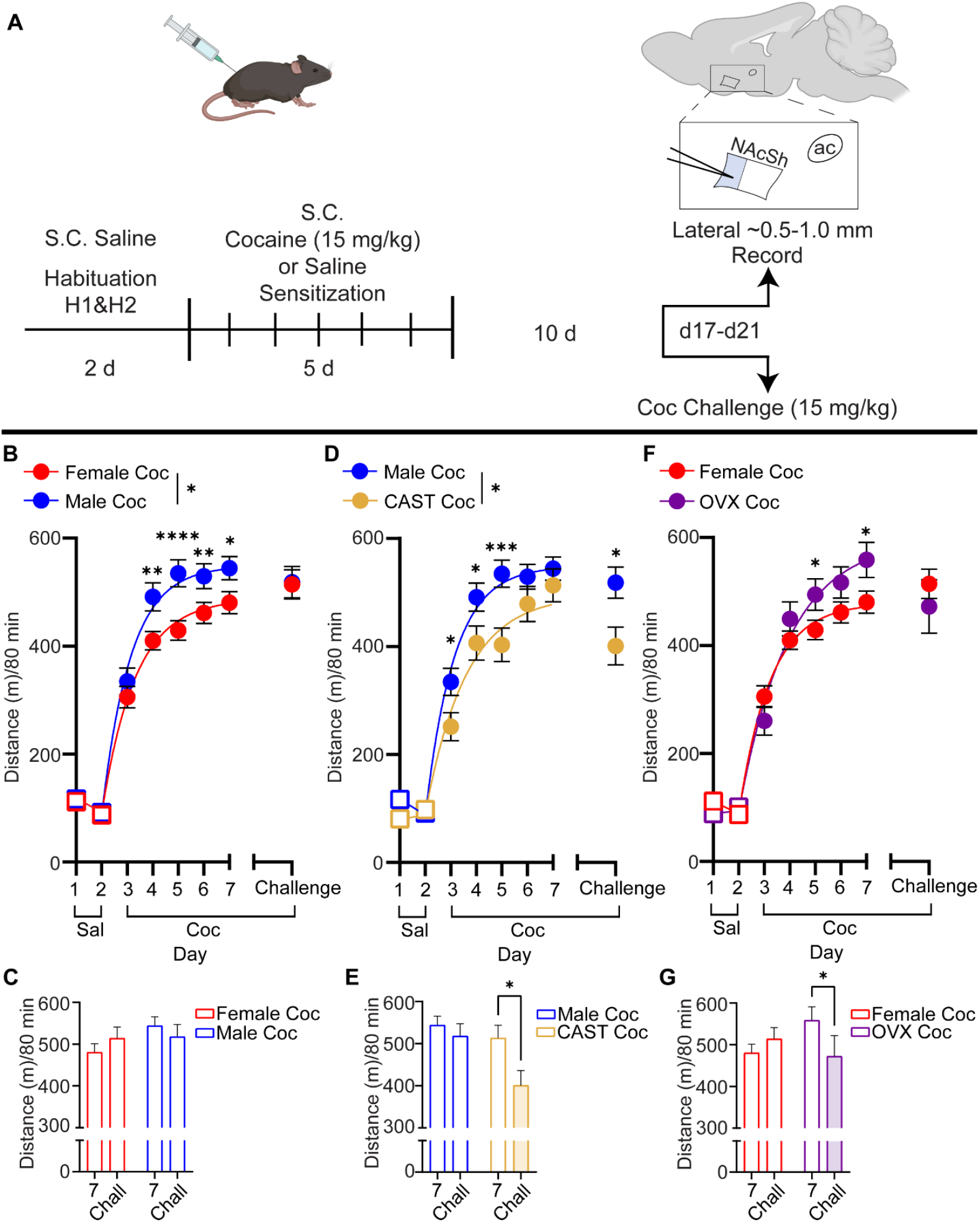
Sex differences and gonadal hormone effect on cocaine psychomotor activity. **A**) Experimental timeline and recording area of NAcSh MSNs. **B)** Male (blue, circles) versus female (red, circles) cocaine sensitization preceded by two days of saline habituation (squares) injections (*p<0.05 two-way ANOVA-Mixed). *p<0.05, **p<0.01, ***p<0.001, ****p<0.0001, Fisher’s LSD. **C)** Total distance traveled comparisons between day 7 cocaine and cocaine challenge after 10-14 days of forced abstinence. **D**) Male (blue, circles) versus CAST male (mustard, circles) cocaine sensitization preceded by two days of saline (squares) habituation injections (*p<0.05 two-way ANOVA-Mixed). *p<0.05, ***p<0.001, Fisher’s LSD. **E)** Total distance traveled comparisons between day 7 and cocaine challenge after 10-14 days of forced abstinence. *p<0.05, two-way ANOVA-RM. **F)** Female (red, circles) versus OVX female (purple, circles) cocaine sensitization preceded by two days of saline (squares) habituation injections. *p<0.05 Fisher’s LSD. **G)** Total distance traveled comparisons between day 7 cocaine and cocaine challenge after 10-14 days of forced abstinence. *p<0.05, two-way ANOVA-RM. Abbreviations: saline (Sal), cocaine (Coc), castrated (CAST), ovariectomized (OVX).

### Circulating gonadal hormones influence cocaine psychomotor activity in both male and female mice

We next wanted to examine the effects of circulating sex hormones on cocaine psychomotor sensitization across animal sex by comparing intact animals to castrated (CAST) male or ovariectomized (OVX) female animals. In males, there was no significant difference during saline habituation in both the intact male group and CAST male group (Fig 1D). In cocaine-treated males and CAST males, cocaine psychomotor activity was greater in the intact males compared to CAST males (Fig 1D; two-way ANOVA-Mixed, F_(1,50)_=4.18, p<0.05). Like intact males, CAST males also sensitized to cocaine (days 3-7, one-way ANOVA-RM, F_(2.73,90.05)_=165.1, p<0.0001). Furthermore, a significant reduction in psychomotor sensitization was apparent in the CAST male group that received a challenge cocaine injection after 10-14 days of abstinence compared to intact males (Fig 1D,E).

In females, OVX increased psychomotor activity compared to intact females (Fig 1F; interaction, two-way ANOVA-Mixed, F_(7,338)_=4.11, p<0.001), with significant differences at days 5 and 7. Like CAST males, OVX females had a significant reduction in cocaine psychomotor sensitization after 10-14 days of abstinence when receiving a challenge injection (Fig 1G, interaction, two-way ANOVA-Mixed, F_(1,22)_=5.06, p<0.05). Overall, loss of gonadal hormones normalized the sex-differences between males and females through a reduction in cocaine psychomotor activity with CAST, and an increase in activity with OVX (Fig S1A,B). OVX increased sensitization such that it equaled intact males (Fig S1C,D), while CAST reduced sensitization such that it equaled intact females (Fig S1E,F).

### Cocaine exposure and abstinence in males reduces NAcSh MSN excitability and requires circulating testicular hormones for cocaine plasticity

To bolster the strength of the electrophysiology throughout this study, we felt it imperative to replicate what has previously been reported in MSNs of the NAcSh from male C57s. This includes a reduction in NAcSh MSN excitability^25^ as well as a potentiation in glutamatergic strength^29, 35^. Thus, the male data was envisioned to serve as a positive control and afford added power to our findings in females, and when investigating the effects of gonadal hormones on cocaine plasticity. In male mice which were exposed to cocaine followed by 10-14 days of abstinence (Fig 1A), we found a reduction in NAcSh MSN excitability (Fig 2A,B, two-way ANOVA-RM, F_(1,39)_=15.7, p<0.001), similar to what has been previously reported^25^. In CAST males, the loss of circulating testicular hormones abolished the apparent reduction in NAcSh MSN excitability induced by cocaine in intact males (Fig 2C,D, two-way ANOVA-RM, F_(1,46)_=1.62, p=0.21). To rule out a floor effect due to CAST in males, we compared the current injection response between intact males and CAST males under saline and cocaine conditions (Fig S2A,D). In the saline treatment groups, there was a mild alteration to the excitability curves, with CAST resulting in differences in firing frequency at 200-220 pA compared to intact males (Fig S2A; two-way ANOVA-RM, treatment x current, F_(8,408)_=4.87, p=0.0001). However this did not result in a floor effect, as comparisons between intact males and CAST males exposed to cocaine still showed a cocaine-induced reduction in NAcSh MSN excitability (140-200 pA) in the intact male group compared to the CAST male group (Fig S2D; two-way ANOVA-RM, treatment x current, F_(8,312_=2.20, p<0.05). Collectively, this suggests that testicular hormones in males are required for cocaine-induced plasticity in MSN excitability of the NAcSh.

**Fig 2.**
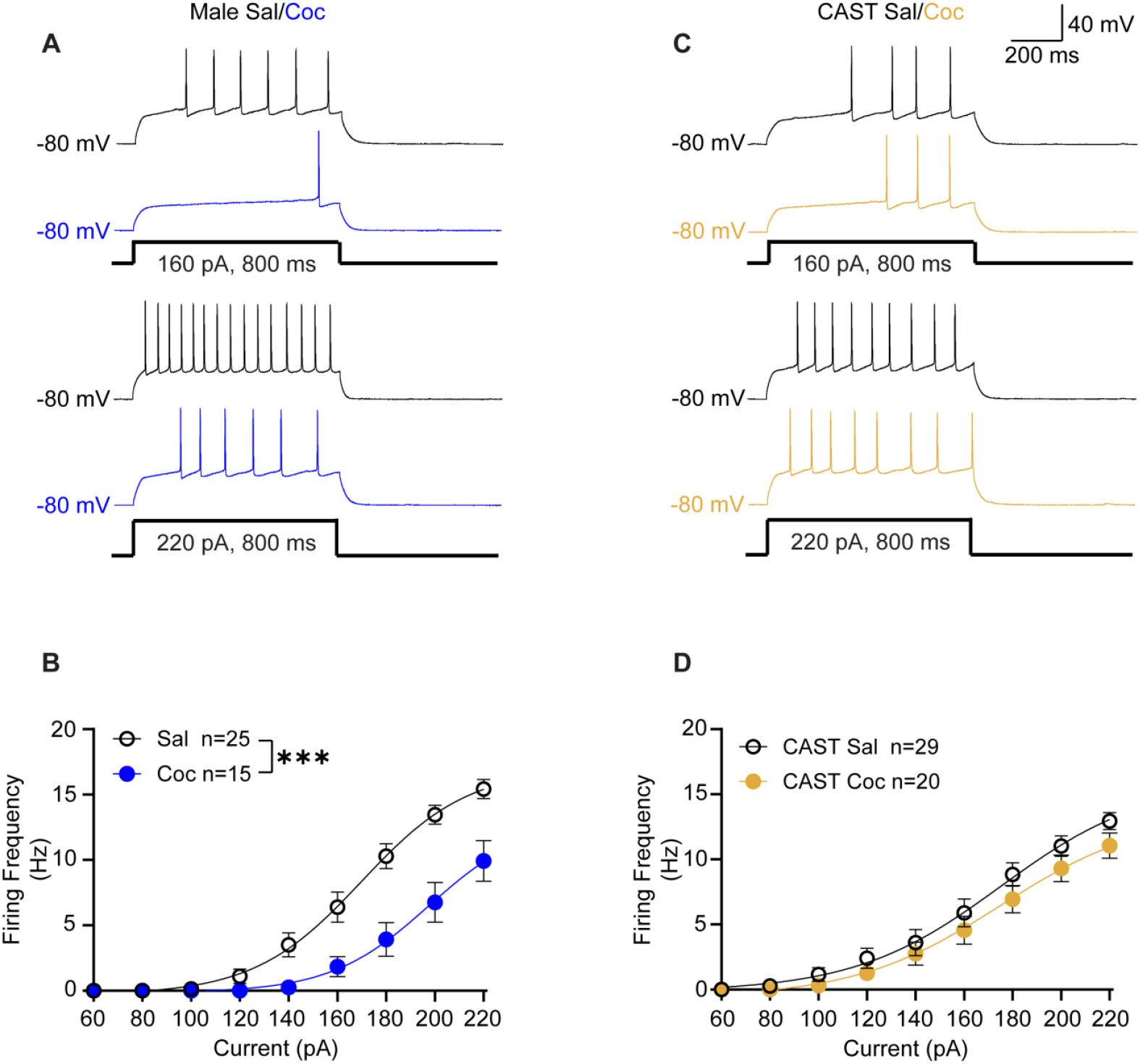
Testicular hormones in male C57s are required for cocaine plasticity in NAcSh MSN excitability. **A)** Representative NAcSh MSN voltage traces from saline treated (black) and cocaine treated (blue) intact male C57s at +160 (top) and +220 (bottom) pA. **B)** Summary data for the current injection response in NAcSh MSNs from saline (black) and cocaine (blue) -treated intact male C57s. ***p<0.001, two-way ANOVA. **C)** Representative NAcSh MSN voltage traces from saline treated (black) and cocaine treated (mustard) castrated male C57s at +160 (top) and +220 (bottom) pA. **D)** Summary data for the current injection response in NAcSh MSNs from saline (black) and cocaine (mustard) -treated castrated male C57s. All neuronal recordings were obtained from n=3-6 mice, number of neurons is reported in the summary data. Abbreviations: Castrated (CAST), saline (Sal), cocaine (Coc).

### Cocaine-induced plasticity in excitability of male MSNs of the NAcSh involves voltage-gated sodium channels and requires testicular hormones

To investigate a potential mechanism for the reduced excitability in male C57 mice following cocaine exposure and abstinence, we performed a ramp injection. This was designed to detect potential differences in voltage threshold to firing an action potential (indicative of alterations to voltage-gated sodium channels), as well as rheobase (the amount of current required to elicit an action potential). In males exposed to cocaine and in abstinence, we detected a depolarizing shift in the voltage threshold to fire an action potential (Fig 3A,B; unpaired t-test, p=0.003), as well as an increased rheobase (Fig 3A,C; unpaired t-test, p<0.0001). Testicular hormones were required for this cocaine-induced alteration, as CAST abolished alterations to voltage threshold to fire an action potential (Fig 3D, E) as well as increased rheobase (Fig 3D, F) between saline and cocaine-treated animals. To investigate whether there were floor effects, we compared voltage threshold to fire an action potential and rheobase in saline-treated intact males and CAST males (Fig S2B, C). There were no differences in alterations to these parameters. When comparing within cocaine treatment in males, intact males did not have a significant difference in voltage threshold to action potential (Fig S2E), although there still was an increased rheobase compared to CAST cocaine-treated males (Fig S2F; unpaired t-test, p<0.001).

**Fig 3.**
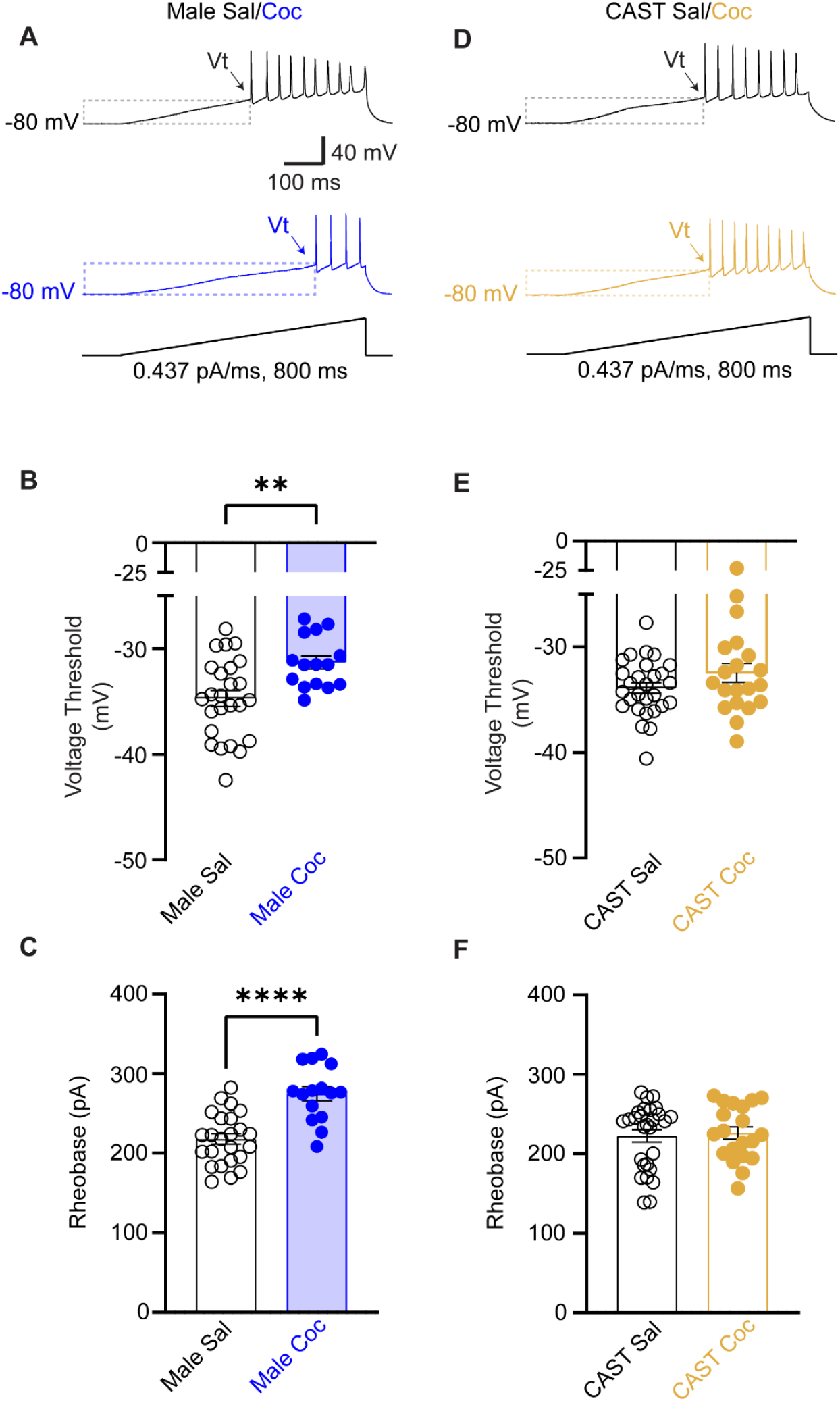
Testicular hormones are required for cocaine-induced alterations to voltage-gated sodium channels. **A)** Representative NAcSh MSN ramp traces from saline-treated (black) and cocaine-treated (blue) intact male C57s. **B)** Summary data for the voltage threshold (Vt) to action potential in NAcSh MSNs from saline (black) and cocaine (blue) -treated intact male C57s. **p<0.01, unpaired t-test **C)** Summary data for rheobase in NAcSh MSNs from saline (black) and cocaine (blue) -treated intact male C57s. ****p<0.0001, unpaired t-test. **D)** Representative NAcSh MSN ramp traces from saline-treated (black) and cocaine-treated (mustard) castrated male C57s. **E)** Summary data for the voltage threshold (Vt) to action potential in NAcSh MSNs from saline (black) and cocaine (mustard) -treated castrated male C57s. **F)** Summary data for rheobase in NAcSh MSNs from saline (black) and cocaine (mustard) -treated intact male C57s. All neuronal recordings were obtained from n=3-6 mice, number of neurons is reported in the summary data. Abbreviations: Castrated (CAST), saline (Sal), cocaine (Coc).

### Cocaine induces an estrous cycle dynamic plasticity in NAcSh MSN excitability and requires ovarian hormones

In females, we first assessed if the estrous cycle affected NAcSh MSN excitability, as ovarian hormones have been shown to alter neuroplasticity in the accumbens core of female rats^36^. As we have previously reported^27, 33^, we were unable to detect any estrous cycle effects in NAcSh MSN excitability in saline-treated female C57s (Fig 4A,B). Next, we examined how cocaine exposure and abstinence impacted NAcSh MSN excitability in estrus and diestrus. In cocaine-exposed females, neuronal excitability recorded during estrus in the abstinence period showed no difference in NAcSh MSN excitability compared to saline-treated females (Fig 4C,D). Conversely, females exposed to cocaine where neuronal excitability was recorded during diestrus in the abstinence period showed reduced NAcSh MSN excitability (Fig 4E,F, two-way ANOVA-RM, F_(1,34)_=9.64, p<0.01). To observe whether ovarian hormones were required for this cocaine plasticity, we performed electrophysiology recordings on saline- and cocaine-treated OVX females. We found that OVX totally abolished any cocaine-induced plasticity to NAcSh MSN excitability (Fig 4G,H), suggesting that ovarian hormones in females are required for NAcSh cocaine plasticity in excitability.

**Fig 4.**
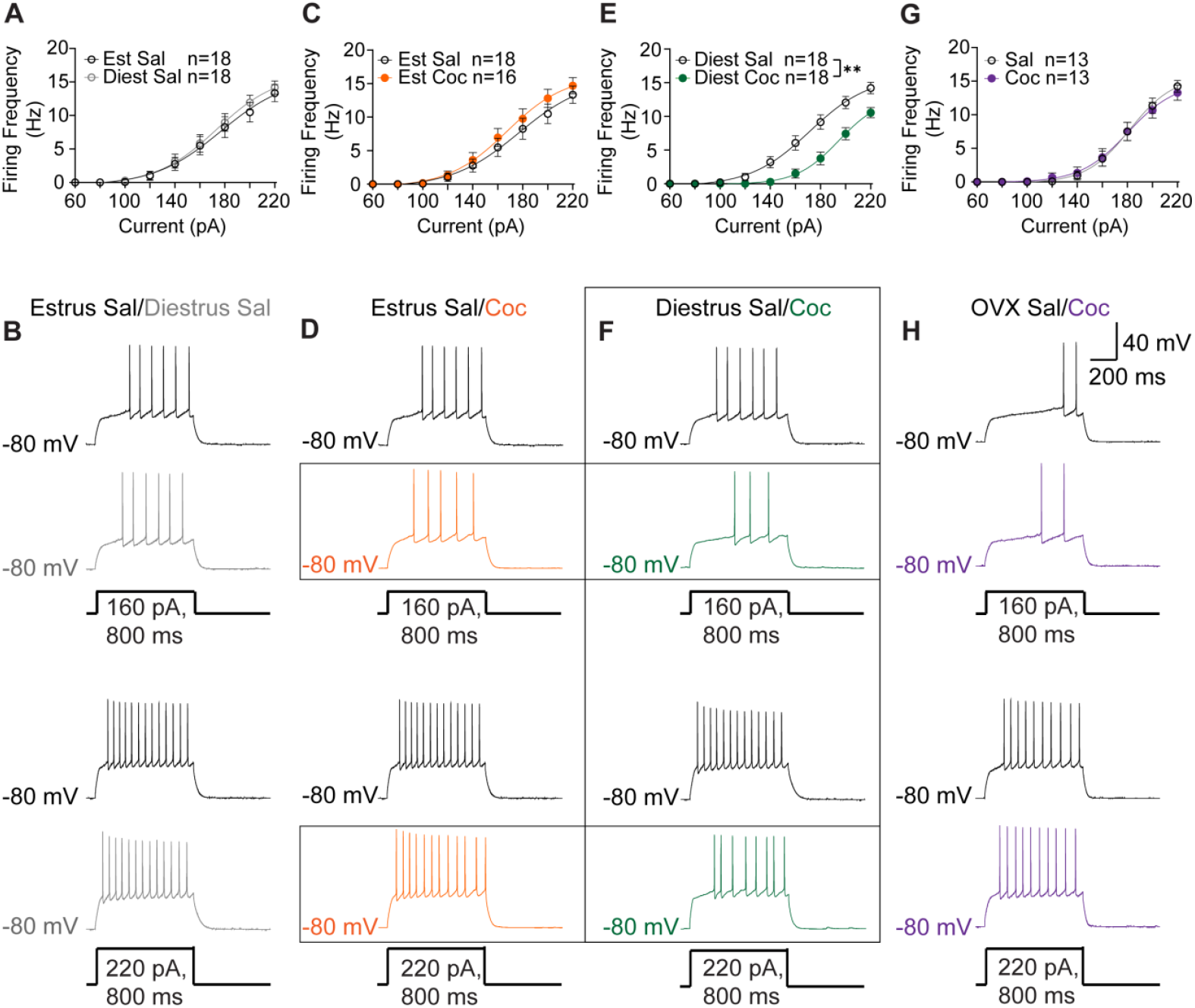
Cocaine induces an estrous cycle-dependent plasticity in NAcSh MSN excitability and requires ovarian hormones. **A)** Summary data for the current injection response in NAcSh MSNs from estrus saline-treated (black) and diestrus saline-treated (grey) intact female C57s. **B)** Representative NAcSh MSN voltage traces from estrus saline-treated (black) and diestrus saline-treated (gray) intact female C57s at +160 (top) and +220 (bottom) pA. **C)** Summary data for the current injection response in NAcSh MSNs from estrus saline-treated (black) and estrus cocaine-treated (orange) intact female C57s. **D)** Representative NAcSh MSN voltage traces from estrus saline-treated (black) and estrus cocaine-treated (orange) intact female C57s at +160 (top) and +220 (bottom) pA. **E)** Summary data for the current injection response in NAcSh MSNs from diestrus saline-treated (black) and diestrus cocaine-treated (green) intact female C57s. **p<0.01, two-way ANOVA. **F)** Representative NAcSh MSN voltage traces from diestrus saline-treated (black) and diestrus cocaine-treated (green) intact female C57s at +160 (top) and +220 (bottom) pA. **G)** Summary data for the current injection response in NAcSh MSNs from OVX saline-treated (black) and OVX cocaine-treated (purple) female C57s. **H)** Representative NAcSh MSN voltage traces from OVX saline-treated (black) and OVX cocaine-treated (purple) female C57s at +160 (top) and +220 (bottom) pA. All neuronal recordings were obtained from n=3-6 mice, number of neurons is reported in the summary data. Abbreviations: Ovariectomy (OVX), saline (Sal), cocaine (coc), estrus (Est), diestrus (Diest).

To compare whether OVX altered inherent neuronal excitability, we compared saline treatment across all female groups. We also compared cocaine treatment across all female groups. In the saline treatment groups, we found no differences in excitability (Fig S3A). However, comparisons in the cocaine treatment groups among females revealed an effect of cocaine between estrus and diestrus (Fig S3D; two-way ANOVA-RM, F_(2,59)_=4.97, p<0.01), as well as a mild effect between estrus and OVX at +160 pA current injection.Again, this reconfirms the effect of the estrous cycle and ovarian hormones intersecting with cocaine plasticity in NAcSh MSN excitability.

### Cocaine-induced plasticity in excitability of female MSNs of the NAcSh involves voltage-gated sodium channels and requires ovarian hormones

In females we also performed a ramp current injection to investigate the potential contribution of voltage gated sodium channels in the cocaine-induced plasticity to NAcSh MSNs. At baseline, there were no estrous cycle effects in voltage threshold to fire an action potential or differences in rheobase (Fig 5A-C). Electrophysiology recordings from cocaine-treated females performed during estrus also had no difference in voltage threshold to fire an action potential or rheobase compared to saline-treated females (Fig 5D-F).

**Fig 5.**
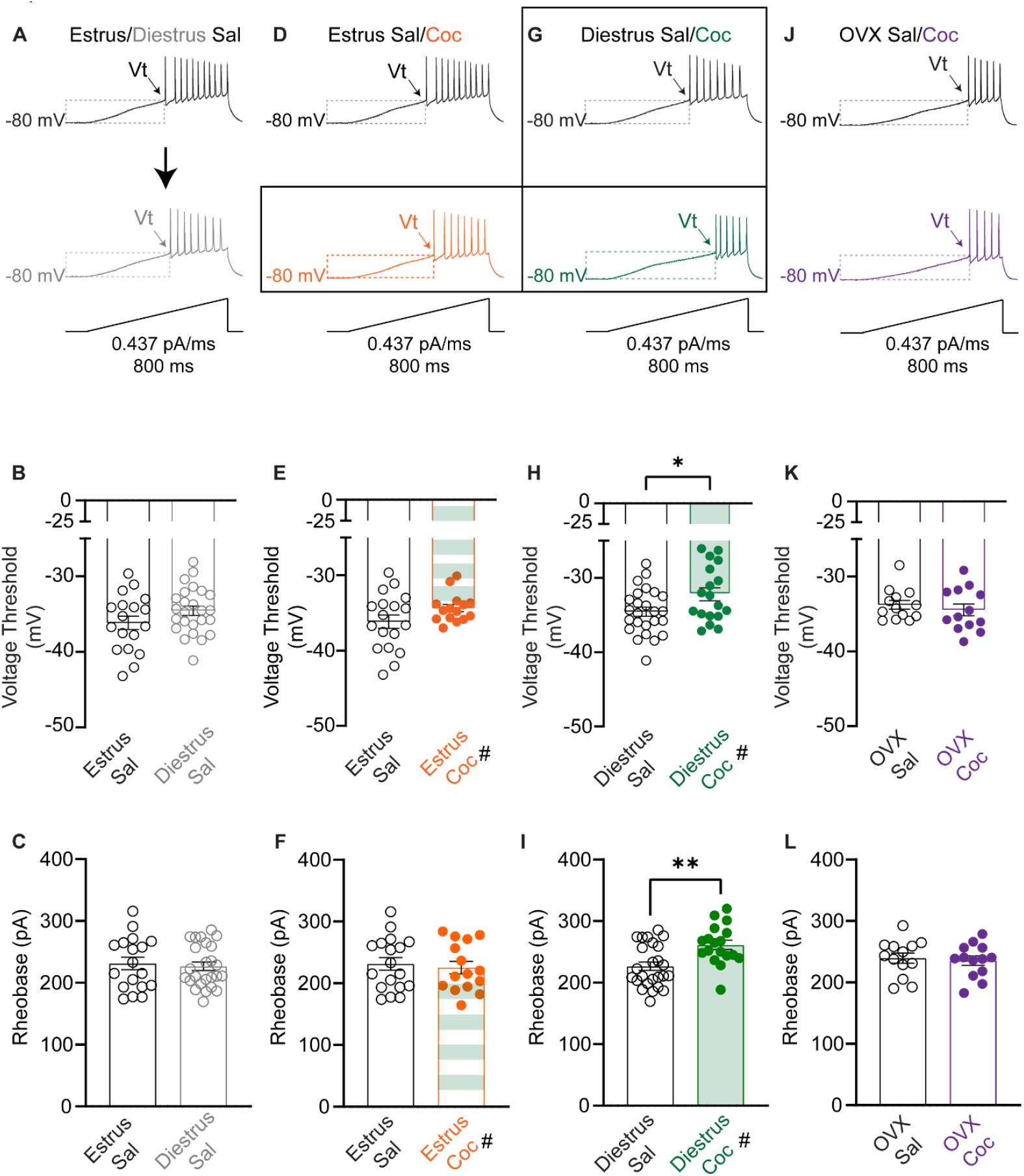
Cocaine induces an estrous cycle dependent plasticity in voltage-gated sodium channels and requires ovarian hormones. **A)** Representative NAcSh MSN ramp traces from estrus saline-treated (black) and diestrus saline-treated (gray) intact female C57s. **B)** Summary data for the voltage threshold (Vt) to action potential in NAcSh MSNs from estrus saline-treated (black) and diestrus saline-treated (gray) intact female C57s. **C)** Summary data for rheobase in estrus saline-treated (black) and diestrus saline-treated (gray) intact female C57s. **D)** Representative NAcSh MSN ramp traces from estrus saline-treated (black) and estrus cocaine-treated (orange) intact female C57s. **E)** Summary data for the voltage threshold (Vt) to action potential in NAcSh MSNs from estrus saline-treated (black) and estrus cocaine-treated (orange) intact female C57s. Note green hatched bars indicate a significant difference between estrus cocaine and diestrus cocaine, #p<0.05. **F)** Summary data for rheobase in estrus saline-treated (black) and estrus cocaine-treated (orange) intact female C57s. Note green hatched bars indicate a significant difference between estrus cocaine and diestrus cocaine, #p<0.05. **G)** Representative NAcSh MSN ramp traces from diestrus saline-treated (black) and diestrus cocaine-treated (green) intact female C57s. **H)** Summary data for the voltage threshold (Vt) to action potential in NAcSh MSNs from diestrus saline-treated (black) and diestrus cocaine-treated (green) intact female C57s. *p<0.05, unpaired t-test. **I)** Summary data for rheobase in diestrus saline-treated (black) and diestrus cocaine-treated (green) intact female C57s. *p<0.05, unpaired t-test. **J)** Representative NAcSh MSN ramp traces from OVX saline-treated (black) and OVX cocaine-treated (purple) female C57s. **K)** Summary data for the voltage threshold (Vt) to action potential in NAcSh MSNs from OVX saline-treated (black) and OVX cocaine-treated (purple) female C57s. **L)** Summary data for rheobase in OVX saline-treated (black) and OVX cocaine-treated (purple) female C57s. All neuronal recordings were obtained from n=3-6 mice. Abbreviations: Ovariectomy (OVX), saline (Sal), cocaine (Coc).

Electrophysiology recordings from cocaine-treated females performed during diestrus revealed a depolarizing shift in voltage threshold to fire an action potential and an increased rheobase compared to saline-treated females (Fig 5G-I; unpaired t-test, p<0.05 and p<0.01, respectively). This finding was also estrous cycle dynamic, as the difference in voltage threshold to fire an action potential and increased rheobase was different between estrus and diestrus cocaine-treated animals, denoted by the hatched bars (Fig 5E,H,F,I; unpaired t-test, p<0.05). OVX totally abolished the effect of cocaine on voltage threshold to fire an action potential and increased rheobase, suggesting that ovarian hormones in females are also required for this effect (Fig 5J-L).

Examining the voltage threshold to action potential and rheobase among all female saline-treated groups, we found that OVX caused a depolarizing shift in voltage threshold to fire an action potential (Fig S3B) and no difference in rheobase (Fig S3C). However, examining voltage threshold to fire an action potential and rheobase among all female cocaine-treated groups revealed a depolarizing shift in voltage threshold to action potential and increased rheobase in diestrus when compared to estrus and OVX females (Fig S3E,F; one-way ANOVA-RM, F_(2,50)_=5.42, p<0.01 and F_(2,50)_=8.15, p<0.001, respectively). This finding suggests an effect of cocaine on voltage-gated sodium channels that are estrous cycle- and ovarian hormone-dependent.

### Gonadectomy produces similar NAcSh MSN excitability in saline and cocaine- and cocaine-treated males and females

To determine whether gonadectomy created any sex-dependent differences in NAcSh MSN excitability, we compared saline-treated CAST males and OVX females. In these two groups, there were mild differences in NAcSh MSN excitability at +140-160 pA (Fig S4A), with no effect on voltage threshold to action potential (Fig S4B), or rheobase (Fig S4C). In cocaine-treated CAST males and OVX females, we detected no difference in NAcSh MSN excitability (Fig S4D), voltage threshold to action potential (Fig S4E), or rheobase (Fig S4F).

### Cocaine potentiates glutamatergic strength in male NAcSh MSNs and partially involves testicular hormones

Cocaine exposure and abstinence potentiate glutamatergic strength in C57 males, reflected by an increase in mEPSC frequencies and amplitudes in MSNs of the NAcSh^35^. Here, we replicated these findings and show that cocaine potentiates NAcSh MSNs glutamatergic strength in males, with an increase in mEPSC frequency and amplitude compared to saline-treated animals (Fig 6A-C; unpaired t-test, p<0.01 and p<0.05, respectively). To understand how testicular hormones influenced glutamatergic plasticity in NAcSh MSNs following cocaine, we also recorded in CAST males. We found that CAST abolished the increase in mEPSC amplitude but not frequency (Fig 6D-F; unpaired t-test, p<0.05). This finding suggests that testicular hormones in males are likely necessary for cocaine-induced postsynaptic AMPAR plasticity, but they do not influence presynaptic glutamate release.

**Fig 6.**
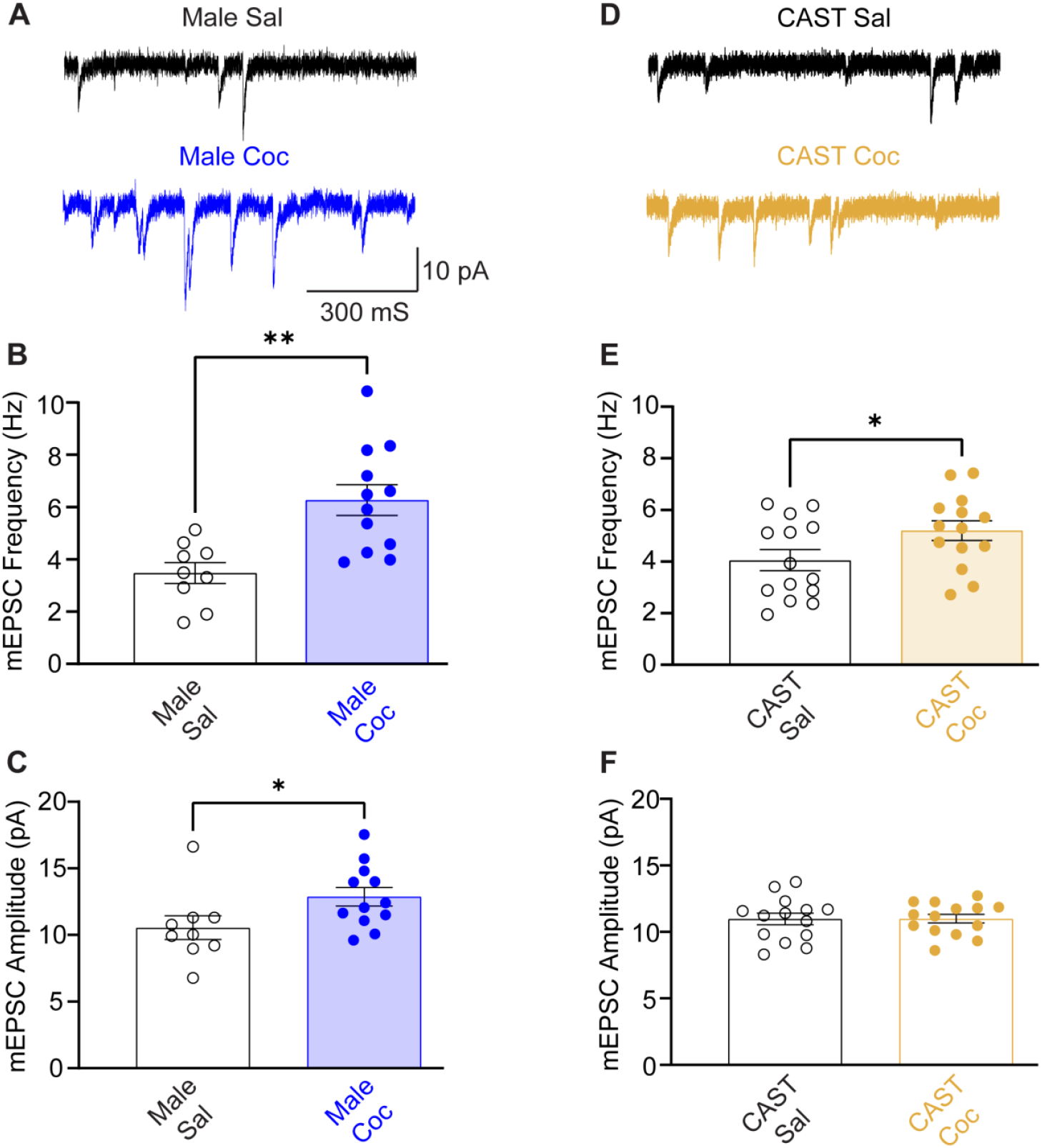
Cocaine increases NAcSh MSN glutamatergic strength and is partially dependent on testicular hormone in males. **A)** Representative NAcSh MSN mEPSC trace from saline-treated (black) and cocaine-treated (blue) intact male C57s. **B)** Summary data for NAcSh MSN mEPSC frequencies from saline (black) and cocaine (blue) -treated intact male C57s. **p<0.01. **C)** Summary data for NAcSh MSNs mEPSC amplitudes from saline (black) and cocaine (blue) -treated intact male C57s. *p<0.05. **D)** Representative NAcSh MSN mEPSC traces from saline-treated (black) and cocaine-treated (mustard) castrated male C57s. **E)** Summary data for NAcSh MSN mEPSC frequencies from saline (black) and cocaine (mustard) - treated castrated male C57s. *p<0.05. **F)** Summary data for NAcSh MSNs mEPSC amplitudes from saline (black) and cocaine (mustard) -treated castrated male C57s. All neuronal recordings were obtained from n=3-6 mice. Abbreviations: Castrated (CAST), saline (Sal), cocaine (Coc).

### Estrous cycle-dependent plasticity in glutamatergic strength in female NAcSh MSNs intersects with cocaine plasticity and requires ovarian hormones

For our mEPSC analysis in females, we first assessed for any baseline estrous cycle plasticity. The estrous cycle has been found to affect mEPSC in the NAc core^36^. Similar to these findings in the core, we also found an estrous cycle-dependent plasticity in NAcSh mESPC frequencies, where there was a reduction in mEPSC frequencies in diestrus compared to estrus (Fig 7A,B; unpaired t-test, p<0.05). There were no differences in mEPSC amplitudes between diestrus and estrus in saline-treated animals (Fig 7A,C). Contrary to the potentiation in glutamatergic strength seen in males following cocaine exposure and abstinence, this same set of conditions in females produced a depotentiation in glutamatergic strength in NAcSh MSNs. There was a reduction in mEPSC frequencies from cocaine-treated females with electrophysiology recordings performed in estrus compared to saline-treatment (Fig 7D,E; unpaired t-test, p<0.05) with no effect on mEPSC amplitudes (Fig 7D,F). Moreover, electrophysiology recordings performed in diestrus in cocaine-treated females revealed a further reduction in mEPSC amplitudes and a trend in reduced mEPSC frequency compared to diestrus saline-treated females (Fig 7G-I; unpaired t-test, p<0.05), suggesting an ovarian hormone and cocaine-induced depotentiation in glutamatergic strength. OVX completely abolished this cocaine-induced and estrous cycle-dependent plasticity (Fig 7J-L), suggesting that ovarian hormones are required for this response, and can modulate both presynaptic glutamate release in addition to postsynaptic AMPAR signaling.

**Fig 7.**
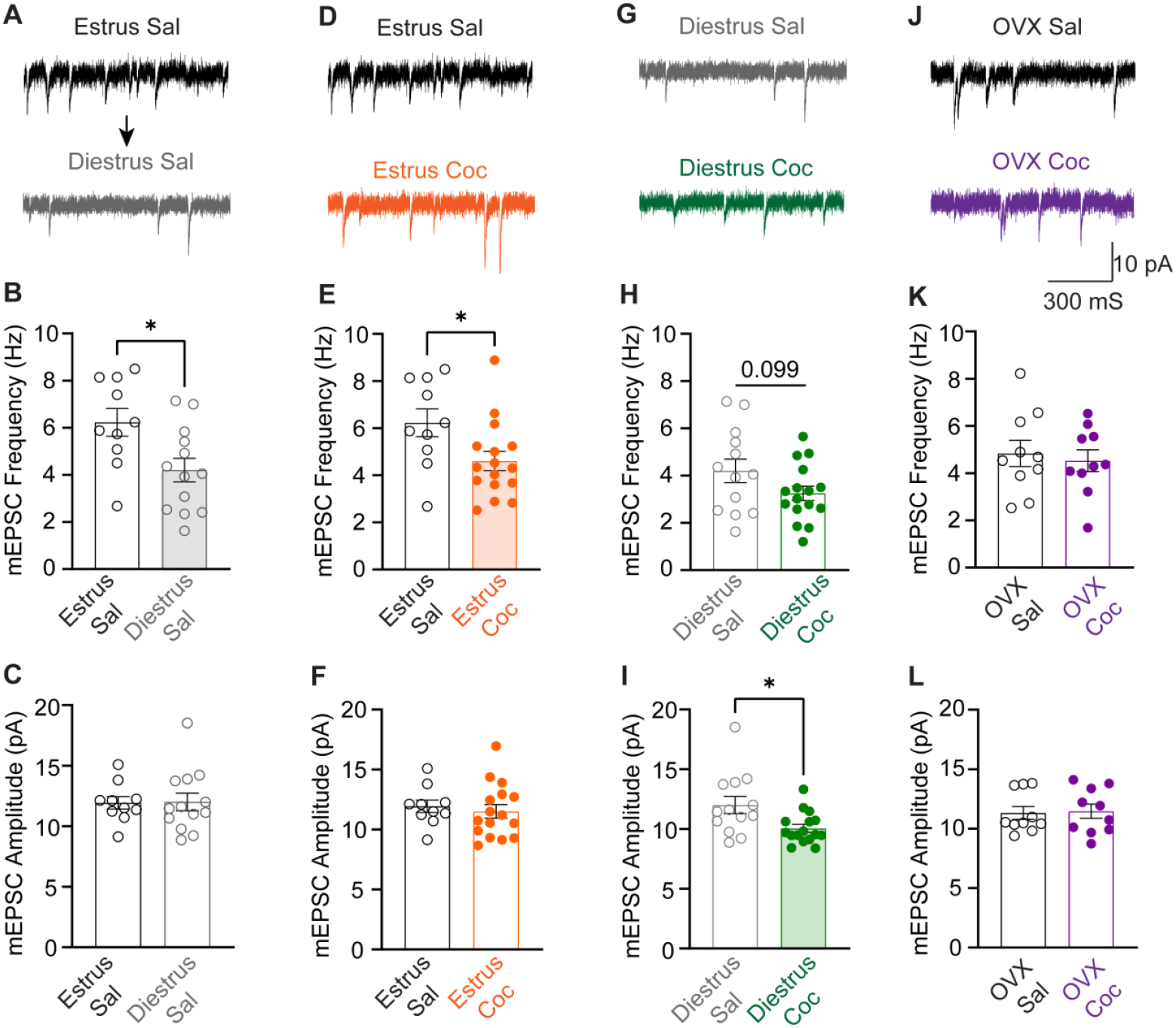
Cocaine decreases NAcSh MSN glutamatergic strength, is estrous cycle- and ovarian hormone-dependent. **A)** Representative NAcSh MSN mEPSC trace from estrus saline-treated (black) and diestrus saline-treated (gray) intact female C57s. **B)** Summary data for NAcSh MSN mEPSC frequencies from estrus saline-treated (black, open circles) and diestrus saline-treated (gray, open circles) intact female C57s. **C)** Summary data for NAcSh MSNs mEPSC amplitudes from estrus saline-treated (black, open circles) and diestrus saline-treated (gray, open circles) intact female C57s. **D)** Representative NAcSh MSN mEPSC traces from estrus saline-treated (black) and estrus cocaine-treated (orange) intact female C57s. **E)** Summary data for NAcSh MSN mEPSC frequencies from estrus saline-treated (black, open circles) and estrus cocaine-treated (orange, closed circles) intact female C57s. *p<0.05. **F)** Summary data for NAcSh MSN mEPSC amplitudes from estrus saline-treated (black, open circles) and estrus cocaine-treated (orange, closed circles). **G)** Representative NAcSh MSN mEPSC trace from diestrus saline-treated (gray) and diestrus cocaine -treated (green) intact female C57s. **H)** Summary data for NAcSh MSN mEPSC frequencies from diestrus saline-treated (gray, open circles) and diestrus cocaine-treated (green, closed circles) intact female C57s. **I)** Summary data for NAcSh MSNs mEPSC amplitudes from diestrus saline-treated (gray, open circles) and diestrus cocaine-treated (green, closed circles) intact female C57s. *p<0.05. **J)** Representative NAcSh MSN mEPSC traces from OVX saline treated (black) and OVX cocaine treated (purple) female C57s. **K)** Summary data for NAcSh MSN mEPSC frequencies from OVX saline treated (black, open circle) and OVX cocaine treated (purple, closed circles) female C57s. **L)** Summary data for NAcSh MSN mEPSC amplitudes from OVX saline treated (black, open circles) and OVX cocaine treated (purple, closed circles) female C57s. All neuronal recordings were obtained from n=3-6 mice. Abbreviations: Ovariectomy (OVX), saline (Sal), cocaine (Coc).

### Gonadal hormones differentially impact mEPSCs

To further understand how glutamatergic strength varied across sex and with gonadal hormones, we compared the mEPSC frequencies and amplitudes under saline conditions and cocaine conditions among all groups. Similar to what Proaño et al. reported in the accumbens core of rats^37^, we saw increased mEPSC frequencies in estrus females compared to diestrus females, intact males, and CAST males in saline-treated animals (Fig S5A). There were no differences in mEPSC amplitudes among saline-treated groups (Fig S5B). Grouped analysis indicated a depotentiation in glutamatergic strength relative to male cocaine-treated animals which was sex specific and estrous cycle dependent (Fig S5C,D). Under cocaine treatment conditions, there were significant reductions in mEPSC frequencies in diestrus females compared to estrus females, intact males, CAST males and OVX females (Fig S5C, one-way ANOVA-RM, F_(4,63)_=6.97, p<0.001). mEPSC amplitudes from cocaine-treated groups were only different between intact males, estrus females, diestrus females, and CAST males, with a reduction relative to intact males in other groups (Fig S5D; one-way ANOVA-RM, F_(4,63)_=4.11, p<0.01).

### Estrous cycle tracking reveals key insights into cocaine-induced plasticity in NAcSh MSNs

Tracking the estrous cycle in females following cocaine exposure and abstinence enhances our understanding of cocaine-induced plasticity. To demonstrate this, we combined our estrus and diestrus saline recordings and compared them to combined estrus and diestrus cocaine recordings for excitability and mEPSCs post hoc. For NAcSh MSN excitability, combined female comparisons revealed no significant differences between saline and cocaine excitability (Fig S6A). For mEPSCs, there was a cocaine-induced depotentiation in glutamatergic strength, reflected in reduced mEPSC frequency (Fig S6B; unpaired t-test, p<0.05) and mEPSC amplitude (Fig S6C; unpaired t-test, p<0.05). This suggests that while a lack of tracking of the estrous cycle still shows effects on glutamatergic strength, it obscures the effects on NAcSh MSN excitability.

### Cocaine abstinence engages divergent NAcSh neurophysiology in males and females

To evaluate whether the absence of estrous cycle tracking contributed to the masking of sex differences in NAcSh MSN neurophysiology during cocaine abstinence, we directly compared males and cycling females as a post hoc comparison. When female cocaine data were pooled and compared to males, we observed significant sex differences in both neuronal excitability and glutamatergic plasticity. In males, NAcSh MSN excitability was reduced relative to pooled female cocaine excitability (Fig 8A, two-way ANOVA-RM, treatment x current, F_(8,376)_=2.34, p<0.05) between 160-220 pA current injection. Similarly, male glutamatergic strength was different during cocaine abstinence compared to pooled females, with an increase in both mEPSC frequency and amplitudes in males compared to females (Fig 8A,B, unpaired t-test, p<0.001 and p<0.01). These findings demonstrate that including sex as a biological variable is important in the context of cocaine research, as the neurobiology of cocaine abstinence is inherently different between males and females, at least within the NAcSh.

**Fig 8.**
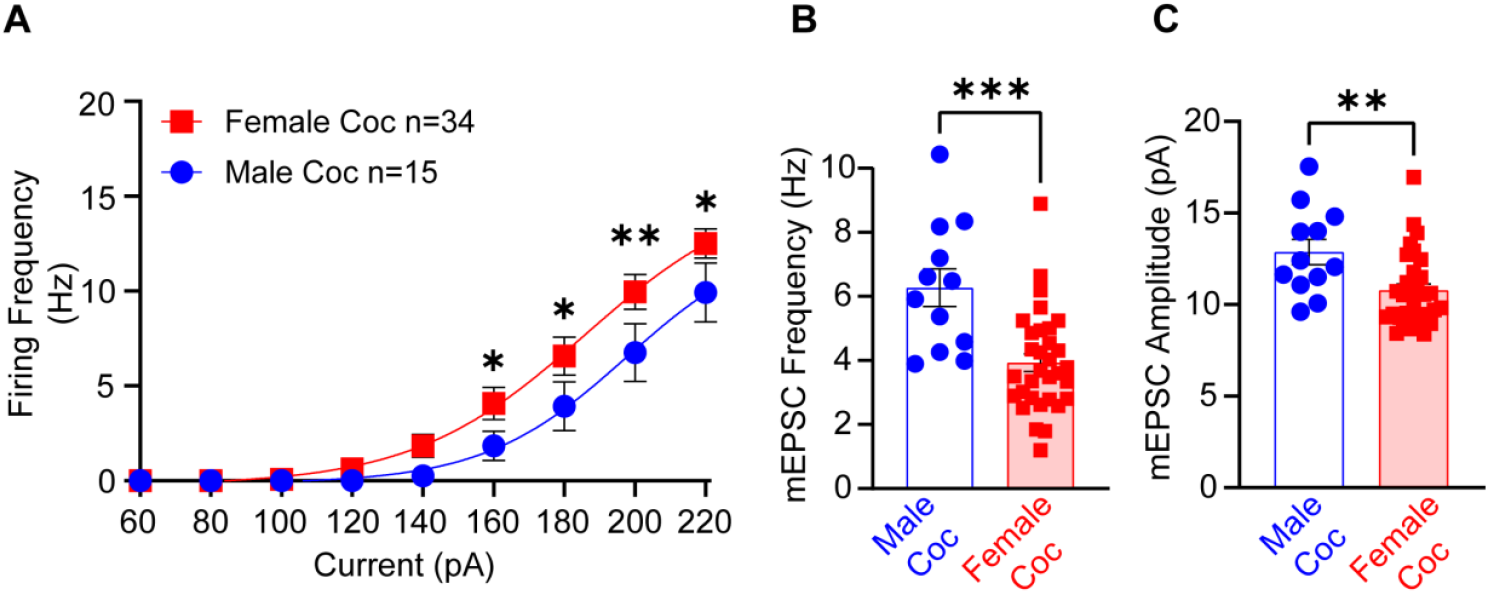
NAcSh MSN neurophysiological comparisons between males and pooled females during cocaine abstinence. **A)** Summary excitability data showing reduced NAcSh MSN excitability from males during cocaine abstinence compared to females, *p<0.05, **p<0.01. **B)** Summary mEPSC frequency data showing an increase in presynaptic glutamate release in males during cocaine abstinence compared to females, ***p<0.001. **C)** Summary mESPC amplitude data showing an increase in postsynaptic AMPAR currents in males during cocaine abstinence compared to females, **p<0.01. All neuronal recordings were obtained from n=3-8 mice, number of neurons is reported in the summary data. Abbreviations: Cocaine (Coc).

### Loss of cocaine-induced neuroplasticity in the NAcSh parallels reduced behavioral plasticity in gonadectomized animals

One secondary question we explored was whether cocaine-induced neuroplasticity in the NAcSh during abstinence is necessary for behavioral plasticity in response to a challenge injection of cocaine. In gonadectomized (GDX) animals, we observed a significant reduction in cocaine-induced neuroplasticity in the NAcSh. Interestingly, this functional loss of neuroplasticity in GDX animals was partially mirrored in behavioral plasticity. While GDX animals did not completely lose cocaine-induced behavioral plasticity, the combined effects of altered neurophysiology in the NAcSh during abstinence and the challenge injection suggest that the NAcSh plays a significant, though partial, role in the psychomotor effects of cocaine sensitization and long-term behavioral changes. Moreover, gonadal hormones in males and ovarian hormones in females are required for these neuroplastic and behavioral adaptations.

## Discussion

Female subjects have been historically understudied in preclinical research. However, their inclusion is important due to physiological, metabolic, and gonadal hormone differences which may influence the brain and behavior differently than in males. The predominant theory in cocaine sensitivity postulates that females are more sensitive to the psychomotor effects of cocaine compared to males^1, 6, 8, 10, 11, 12^. These studies were however, primarily conducted in rats. When investigating the contribution of gonadal hormones in these cocaine responses, OVX has been shown to result in a reduction in cocaine-induced locomotion compared to intact females^38, 39^, while CAST males have an enhanced cocaine response^39^. Thus, the sex differences in cocaine sensitization in rats can be narrowed via GDX. In our hands, the C57 strain has a totally flipped cocaine behavioral phenotype. In C57s, we replicated our previous findings that males show a stronger cocaine psychomotor response than females^21^. Additionally, we now show that OVX enhances cocaine psychomotor effects, while CAST reduces it. In effect, the sex gap was again narrowed, but in an opposite way that has been shown in rats. While we are not entirely sure why the C57s have a reversed behavioral phenotype, genetic background^21, 40, 41, 42, 43, 44, 45, 46, 47^, and potentially species specific (rats vs mice)^47, 48^ may be a contributing factor. Future work is needed to discover the molecular mechanisms that mediate these species differences and how they differ or are similar to humans.

Interestingly, when we investigated how abstinence (10-14 days) altered cocaine behavioral plasticity, we found a narrowing of the sex gap such that males and females had equal responses to a cocaine challenge injection. So, while the initial 5-day sequential cocaine exposure was blunted in females compared to males, the abstinence period seemed to contribute to a building behavioral plasticity in females. This is consistent with incubation of craving responses reported in rats where withdrawal time increased attempts for cocaine^49, 50, 51^. Surprisingly, both OVX and CAST in the C57s drastically reduced this cocaine behavioral plasticity, suggesting that gonadal hormones were required for the sustained cocaine sensitization following abstinence-a finding that to the best of our knowledge has not been reported.

Another arm of our study was to investigate how cocaine exposure influenced neuroplasticity in MSNs of the NAcSh during abstinence. Within the C57 strain, these cocaine-induced alterations to neurophysiology in MSNs of the NAcSh have only been reported in males. This has led to a knowledge gap in how cocaine exposure and abstinence alters female NAcSh neuroplasticity, with treatment strategies relying heavily on male electrophysiology as key predictors of cocaine relapse mechanisms^29, 35, 52, 53, 54^. Because the estrous cycle induces its own neuroplasticity, we anticipated cocaine exposure may intersect with estrous cycle neuroplasticity and obscure potential findings if the estrous cycle was not tracked^22, 55, 56^. Thus, we made recordings in estrus and diestrus, two points during the estrous cycle where the maximum difference in neuroplasticity has been observed^22^. Remarkably, we found that cocaine induced an estrous cycle dynamic effect on NAcSh MSN excitability, where excitability was reduced in diestrus relative to estrus.

If female data was pooled, it obscured differences between saline and cocaine treatment. We also replicated a cocaine-induced reduction in male NAcSh MSN excitability^25^. As a novel addition, we performed excitability recordings in OVX and CAST mice-key neurophysiological parameters which have yet to be reported in any animal model but remains a crucial component to understanding the effects of gonadal hormones on cocaine behavior. Unexpectedly, GDX abolished any cocaine-induced neuroplasticity to MSN excitability in the NAcSh from both male and female mice, at least for long-term cocaine plasticity. Whether GDX also affects short-term cocaine abstinence plasticity remains to be determined but is a necessary follow-up study.

For contributing mechanisms in excitability, the involvement of voltage-gated sodium channels in altering NAcSh MSN excitability following cocaine exposure and abstinence was not totally surprising. Cocaine, as part of a broader class of local anesthetic drugs such as, lidocaine, benzocaine, and procaine, blocks nerve impulses by blocking voltage-gated sodium channels^57^. Indeed, Zhang et al have shown that cocaine exposure and abstinence produce reduced neuronal excitability in the accumbens via reduced whole-cell sodium currents, indicative of alterations to voltage gated sodium channels^58^. Lidocaine failed to replicate the effect of cocaine, suggesting the mechanism was not via local anesthetic properties and more closely aligned with a dopamine-induced alteration in voltage-gated sodium channels^58^. Cocaine-induced dopaminergic dynamics have been well documented by sex in the C57s by Calipari et al, and we speculate that the ventral tegmental area to NAc circuit in the C57s likely influences NAcSh MSN excitability via a dopamine-VGSC mechanism. Alternatively, cocaine-induced epigenetic modification of VGSCs may also underlie this effect^59^.

Similar to the unexpected effects of cocaine on female NAcSh MSN excitability, the effects of cocaine on glutamatergic strength in this brain region were also different in females. Whereas cocaine potentiated glutamatergic strength in males, it was depotentiated in females and was estrous cycle dynamic. OVX completely abolished any effects of cocaine on altering glutamatergic strength, again implicating a requirement of ovarian hormones in females for this effect. CAST in males abolished the postsynaptic AMPAR plasticity but spared the cocaine-induced enhancement in presynaptic glutamatergic plasticity. In C57 males, potentiation in glutamatergic strength is a hallmark of cocaine exposure and abstinence^29, 60^, with relapse or cocaine re-exposure depotentiating glutamatergic strength in MSNs of the NAcSh^29, 35, 52^. This glutamatergic mechanism has been proposed as a major driver of cocaine relapse^52, 53, 61^. Given that pharmacological and optogenetic manipulation of glutamatergic strength via AMPAR-dependent plasticity facilitated cocaine-primed reinstatement^52^, our reported effect of CAST on cocaine AMPAR plasticity in male C57s suggests hormone modulation may be an effective tool for combating cocaine relapse in males.

Likewise, using the glutamatergic mechanism for cocaine reinstatement generated in C57 males^52^ but now applying it to females, cocaine exposure in estrus would be anticipated to depotentiate glutamatergic strength and may account for enhanced cocaine CPP in estrus as reported by Calipari et al^7^. Similarly, hormonal depotentiation from estrus transitioning to diestrus during cocaine abstinence may also facilitate estrous cycle-specific sensitivities. The behavioral parallel of cocaine challenge injections with the electrophysiology in this timeframe strongly supports this hypothesis. Future studies exploring the causality of glutamatergic strength in the NAcSh and cocaine relapse in C57s between sexes with the inclusion of CAST and OVX animals seem warranted and necessary to discern sex- and hormone-specific contributions in these behavioral and neurophysiological effects. Interestingly, hormonal modulation has been suggested as a means to treat cocaine use disorder^62^, and our findings here support that notion.

As a final point, in females, the effects of cocaine abstinence on MSN excitability, VGSC function, and glutamatergic strength were dependent on the estrous cycle, with a majority of changes emerging only after cocaine exposure. These findings suggest that motivated behaviors in females post-cocaine exposure may differ fundamentally from those observed in drug-naive animals. Such altered behaviors could include, but are not limited to reproductive, feeding, and social behaviors. Whether similar alterations in motivation influence drug responses in the reverse direction, for example, prior motivated states affecting drug effects, remains an open question. Nonetheless, our data provide a foundation for understanding how drug-induced adaptations within the reward circuitry may shape future motivated behaviors in females.

In conclusion, this study challenges several longstanding hypotheses about cocaine-related behavior and abstinence-induced plasticity in the NAcSh. Our findings suggest that biological sex, circulating gonadal hormones, and other individual factors may critically influence vulnerability to developing cocaine use disorder. Understanding converging themes across diverse rodent models and genetic backgrounds will likely yield the most robust targeting for treatment in humans. Finally, our findings emphasize that the neuropharmacological actions of cocaine differ fundamentally between the sexes, reinforcing the importance of studying both sexes in preclinical and clinical research^63^.

## Acknowledgments

We would like to thank Brian A. Chapp, Dr. Timothy W. Chapp, Dr. Scott M. Chapp, Dr. Robert Meisel, and Dr. Hannah M. McMullan for their proofreading and suggestions. We would also like to thank the UMN Mouse Behavior Core for their equipment access and technical support. Certain components of the figures were created with biorender.com. A version of this manuscript has been deposited to bioRxiv.

## Author Contributions

A.D.C, C.A.N, C-M.H.P, and P.P.J performed experiments; A.D.C and C.A.N analyzed data; A.D.C, C.A.N, P.G.M, M.J.T, prepared figures; A.D.C, C.A.N, A.R.C, E.B.L., Y.A.C, P.G.M, and M.J.T drafted manuscript; A.D.C, C.A.N, A.R.C, E.B.L, Y.A.C., P.G.M and M.J.T interpreted results of experiment; A.D.C, C.A.N, C-M.H.P, P.P.J, A.R.C, E.B.L., Y.A-C. P.G.M and M.J.T edited and revised the manuscript.

## Graphical summary

Effect of sex, and gonadal hormones on cocaine psychomotor activity and NAcSh MSN neurophysiology during cocaine abstinence in C57BL/6J mice. Arrows between estrus and diestrus denote estrous cycle-dependent plasticity.

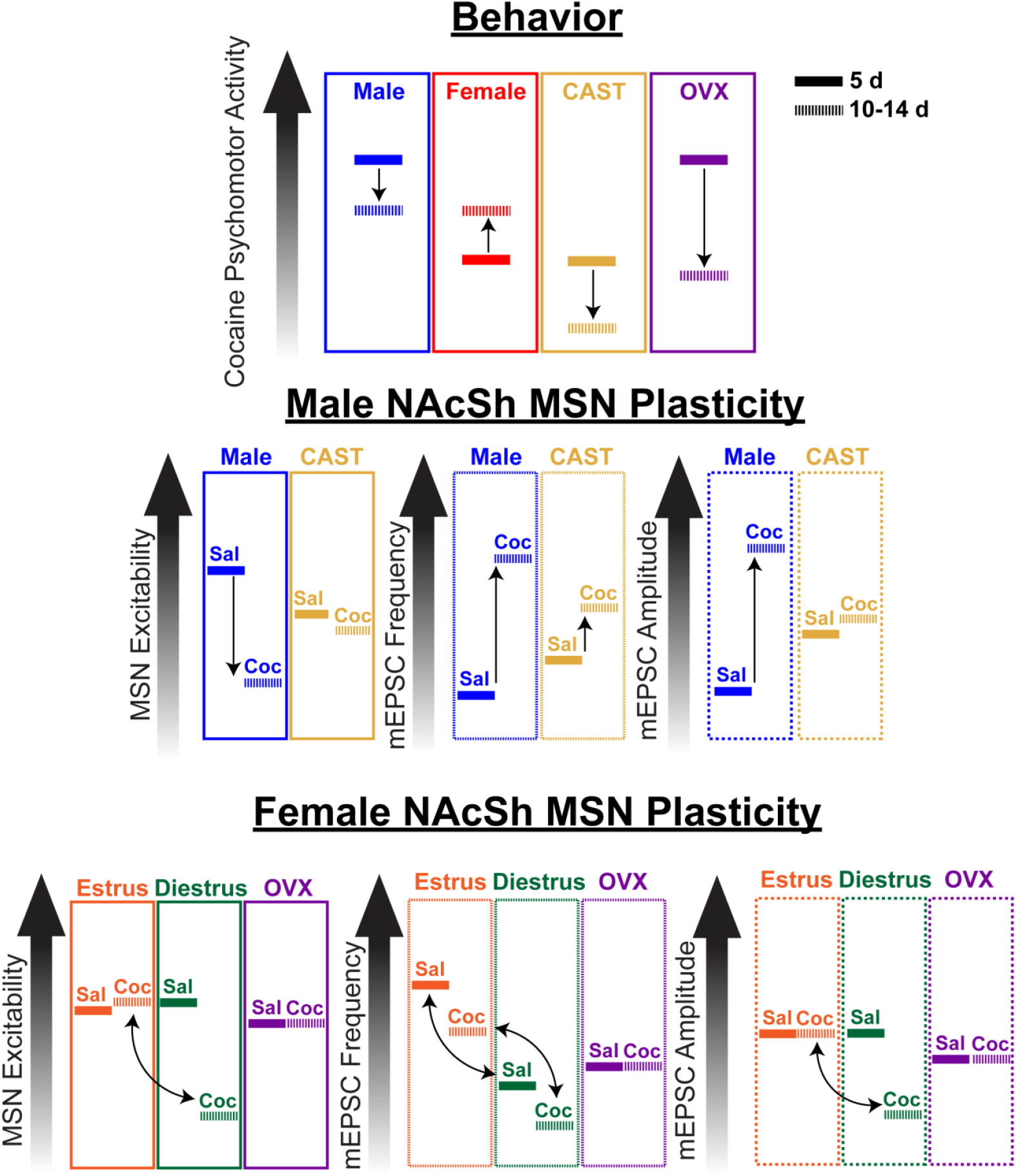

## Supplementary Materials

### Supplemental Figures

**Fig S1.**
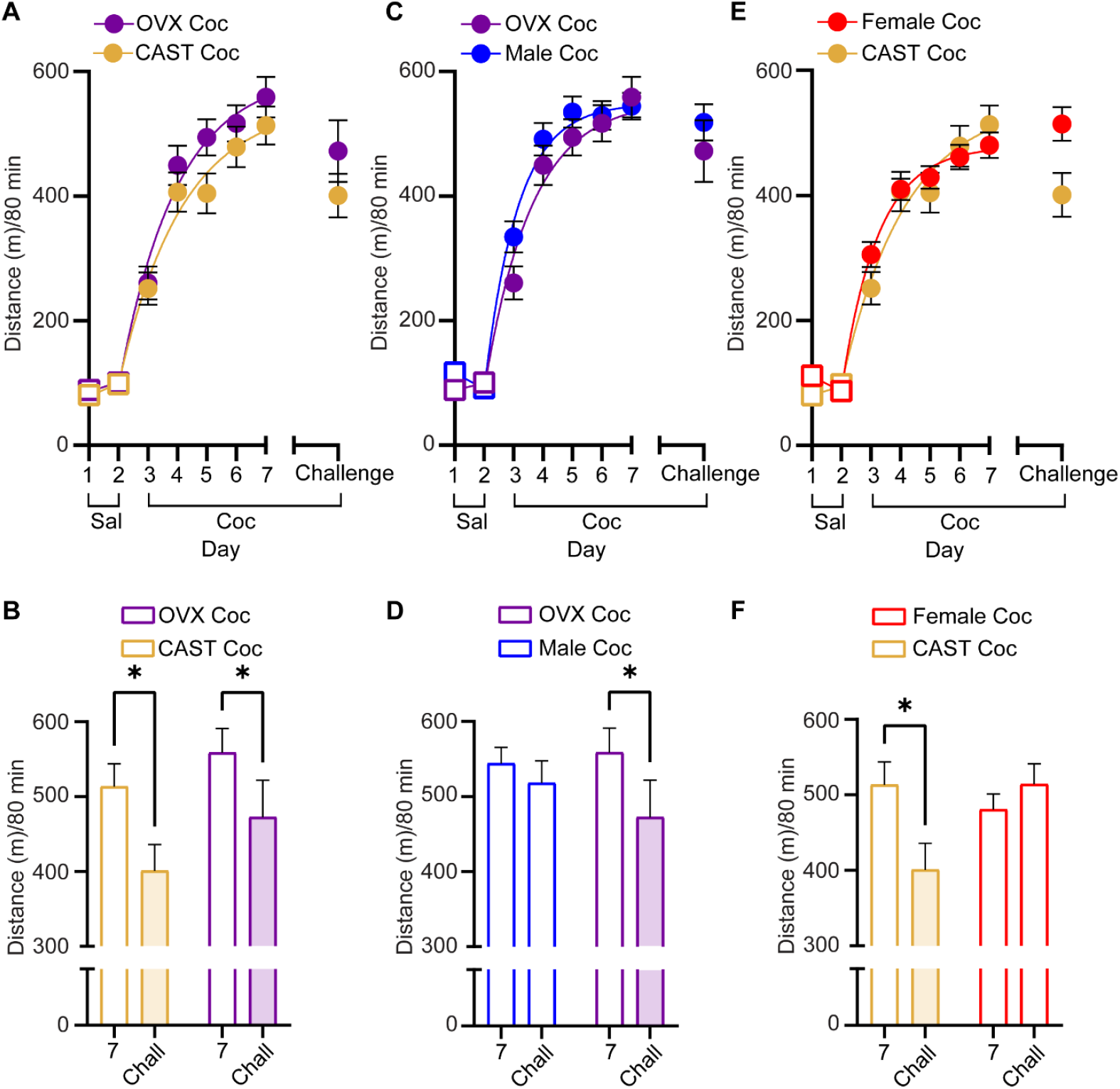
Castration and ovariectomy normalize sex differences in cocaine psychomotor activity and reduce psychomotor plasticity. **A)** Summary cocaine psychomotor sensitization data for comparisons between CAST males (mustard) and OVX females (purple). **B)** Summary data for cocaine psychomotor plasticity between day 7 and challenge cocaine (after 10-14 days abstinence). *p<0.05, two-way ANOVA-Mixed. **C)** Summary cocaine psychomotor sensitization data for comparisons between males (blue) and OVX females (purple). **D)** Summary data for cocaine psychomotor plasticity between day 7 and challenge cocaine (after 10-14 days abstinence). *p<0.05, two-way ANOVA-Mixed. **E)** Summary cocaine psychomotor sensitization data for comparisons between CAST males (mustard) and females (red). **F)** Summary data for cocaine psychomotor plasticity between day 7 and challenge cocaine (after 10-14 days abstinence). *p<0.05, two-way ANOVA-Mixed. Open squares (saline), closed circles (cocaine). Abbreviations: CAST (castration), OVX (ovariectomy), Chall (challenge).

**Fig S2.**
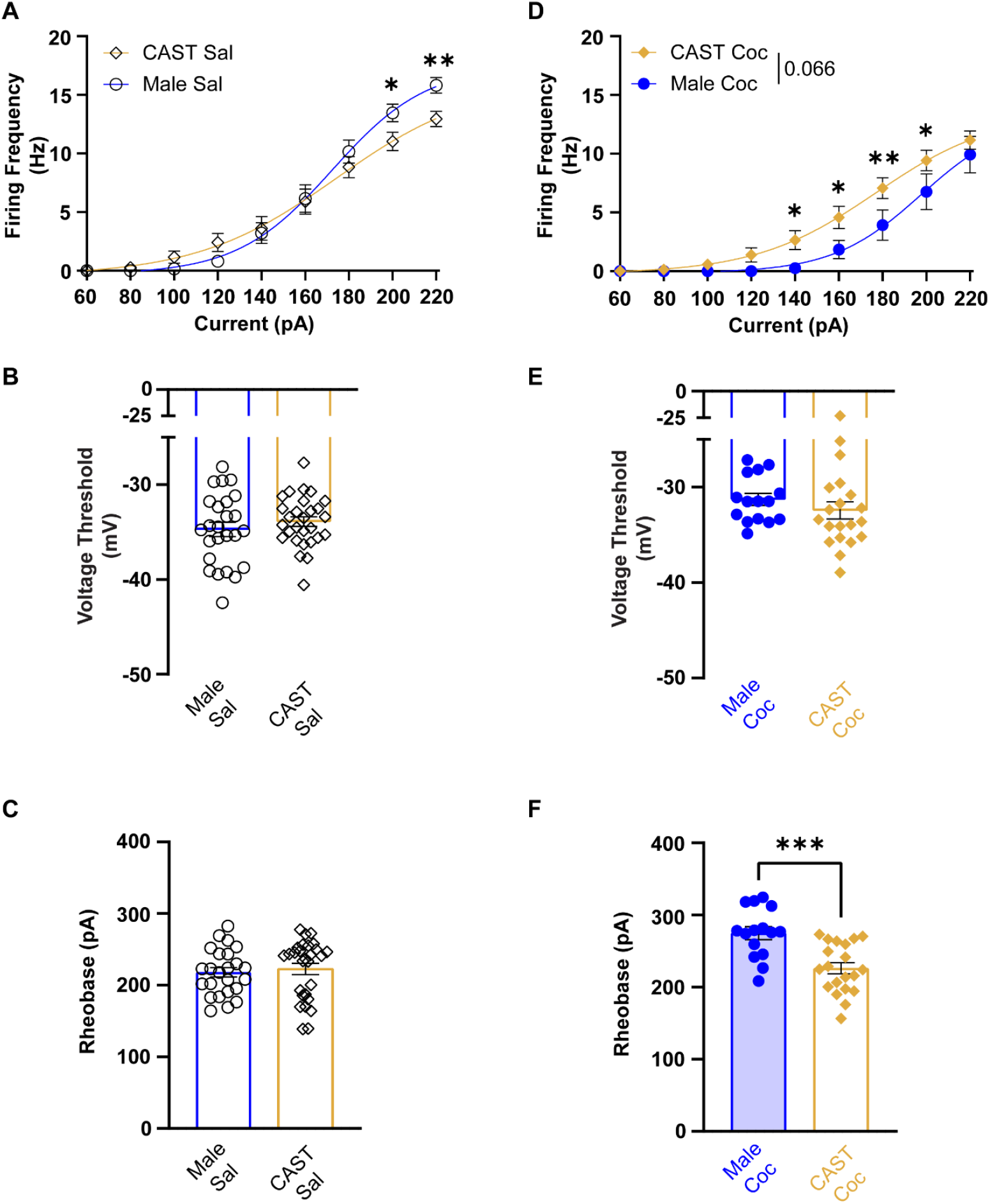
Effect of male gonadal hormones on NAcSh MSN excitability. **A)** Summary data showing the current injection response from castrated males (mustard, open diamonds) and intact male (blue, open circles) saline-treated animals, *p<0.05. **B)** Summary data for voltage threshold (Vt) to firing an action potential (AP) between intact male saline (blue, open circles) and castrated male (mustard, open diamonds) saline-treated animals. **C)** Summary data for rheobase between intact male (blue, open circles) and castrated male (mustard, open diamonds) saline-treated animals. **D)** Summary data showing the current injection response from castrated male (mustard, solid diamonds) and intact male (blue, solid circles) cocaine-treated animals, *p<0.05, **p<0.01. **E)** Summary data for voltage threshold (Vt) to firing an action potential (AP) between intact male (blue, solid circles) and castrated male (mustard, solid diamonds) cocaine-treated animals. **F)** Summary data for rheobase between intact male (blue, solid circles) and castrated male (mustard, solid diamonds) cocaine-treated animals, ***p<0.001. All neuronal recordings were obtained from n=3-6 mice. Abbreviations: saline (Sal), cocaine (Coc), Castrated (CAST).

**Fig S3.**
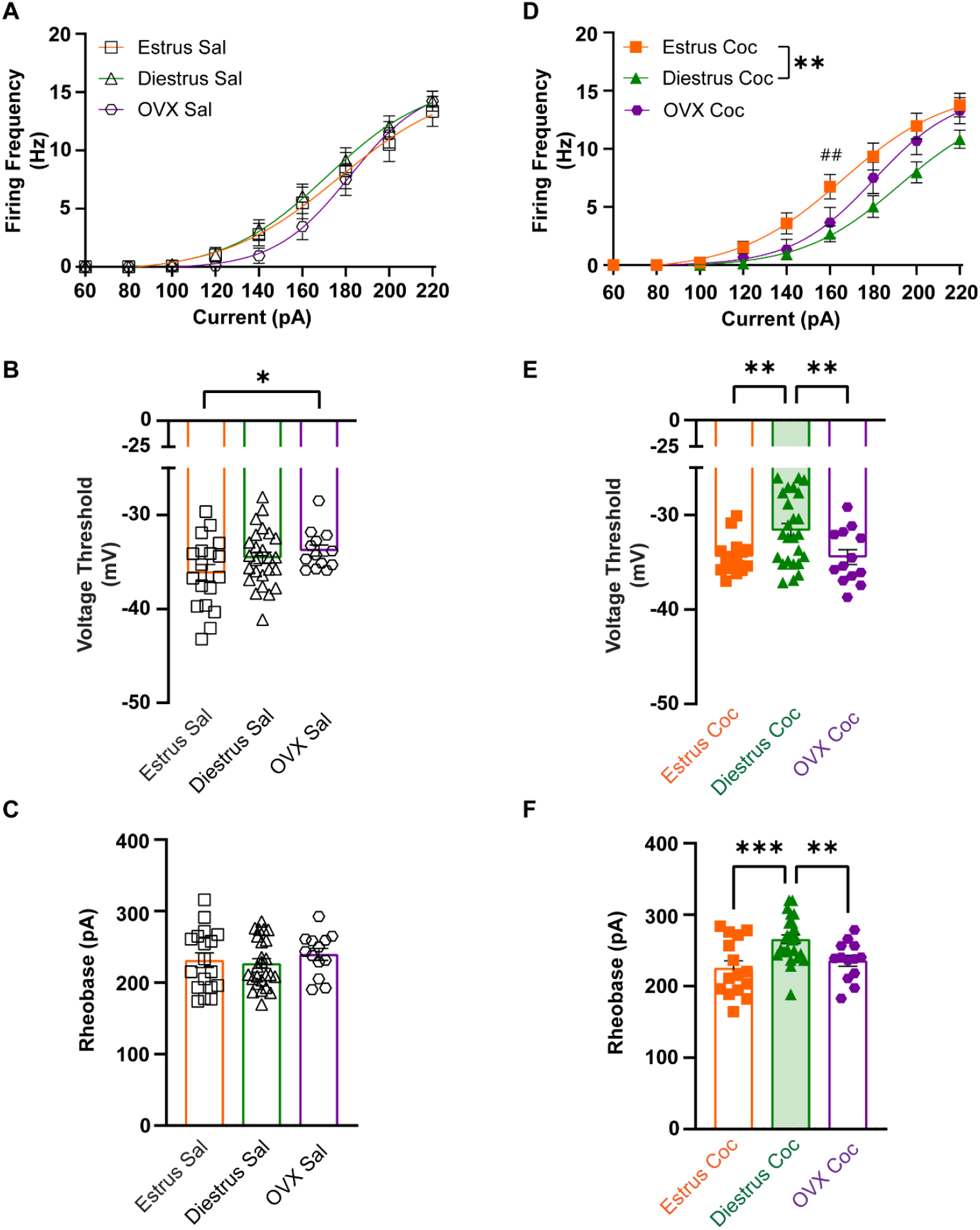
Ovarian hormones influence NAcSh MSN excitability. **A)** Summary data showing the current injection response from estrus female (orange, open square), diestrus female (green, open triangles), and ovariectomized female (purple, open hexagon) saline-treated animals. **B)** Summary data for voltage threshold (Vt) to firing an action potential (AP) from estrus female (orange, open square), diestrus female (green, open triangles), and ovariectomized female (purple, open hexagon) saline-treated animals **C)** Summary data for rheobase from estrus female (orange, open square), diestrus female (green, open triangles), and ovariectomized female (purple, open hexagon) saline-treated animals. **D)** Summary data showing the current injection from estrus female (orange, solid square), diestrus female (green, solid triangles), and ovariectomized female (purple, solid hexagon) cocaine-treated animals, **p<0.01, ## p<0.01 Estrus Coc vs OVX Coc. **E)** Summary data for voltage threshold (Vt) to firing an action potential (AP) from estrus female (orange, solid square), diestrus female (green, solid triangles), and ovariectomized female (purple, solid hexagon) cocaine-treated animals, **p<0.01. **F)** Summary data for rheobase from estrus female (orange, solid square), diestrus female (green, solid triangles), and ovariectomized female (purple, solid hexagon) cocaine-treated animals, **p<0.01, ***p<0.001. All neuronal recordings were obtained from n=3-6 mice. Abbreviations: saline (Sal), cocaine (Coc), ovariectomized (OVX).

**Fig S4.**
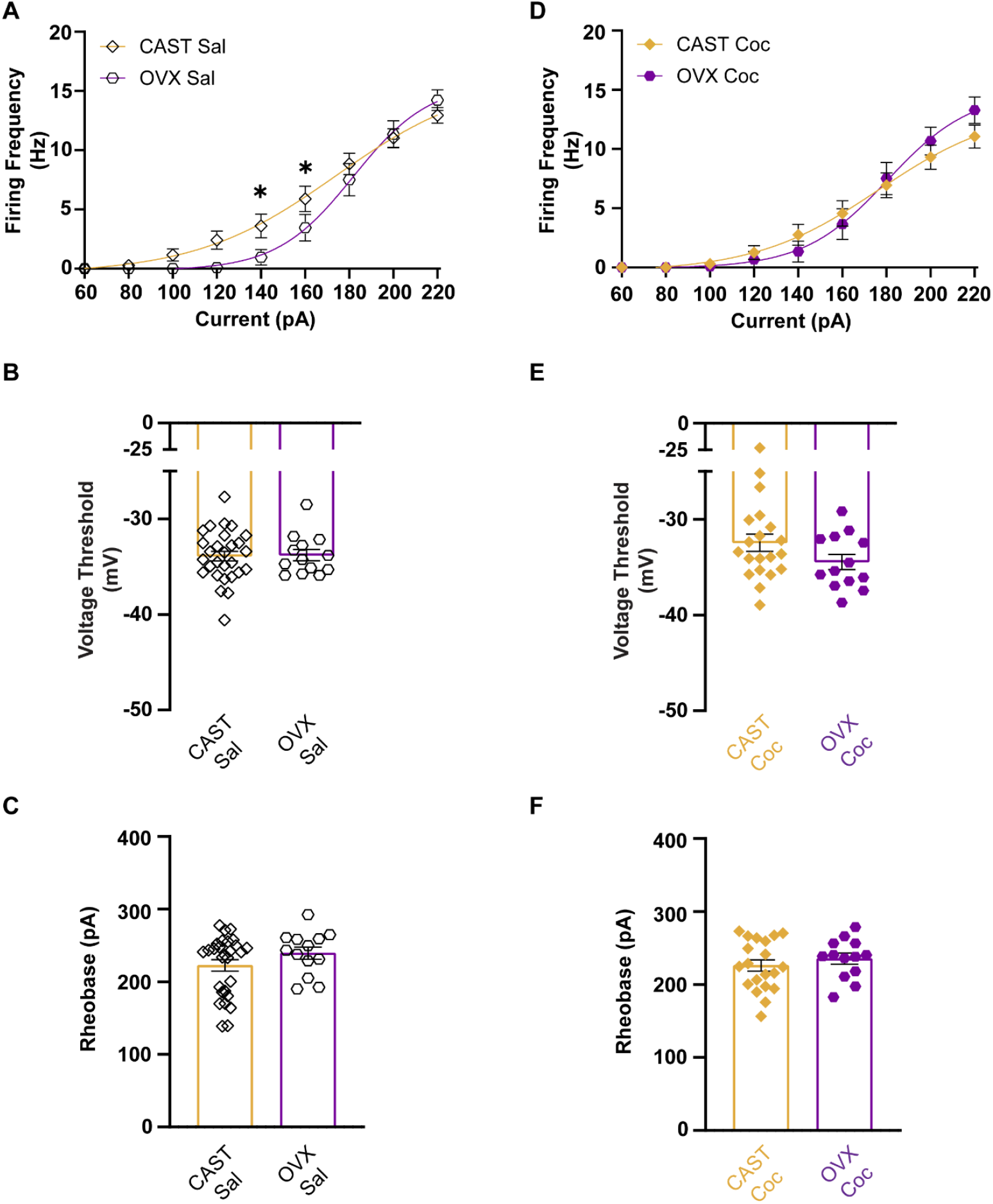
Comparison between castration and ovariectomy on NAcSh MSN excitability between sexes. **A)** Summary data showing the current injection response from castrated males (mustard, open diamonds) and ovariectomized female (purple, open hexagon) saline-treated animals. **B)** Summary data for voltage threshold (Vt) to firing an action potential (AP) between castrated males (mustard, open diamonds) and ovariectomized female (purple, open hexagon) saline-treated animals. **C)** Summary data for rheobase between castrated males (mustard, open diamonds) and ovariectomized female (purple, open hexagon) saline-treated animals. **D)** Summary data showing the current injection response between castrated males (mustard, solid diamonds) and ovariectomized female (purple, solid hexagon) cocaine treated animals. **E)** Summary data for voltage threshold (Vt) to firing an action potential (AP) between castrated males (mustard, solid diamonds) and ovariectomized female (purple, solid hexagon) cocaine-treated animals. **F)** Summary data for rheobase between castrated males (mustard, solid diamonds) and ovariectomized female (purple, solid hexagon) cocaine-treated animals. All neuronal recordings were obtained from n=3-6 mice. Abbreviations: saline (Sal), cocaine (Coc), castrated (CAST), ovariectomized (OVX).

**Fig S5.**
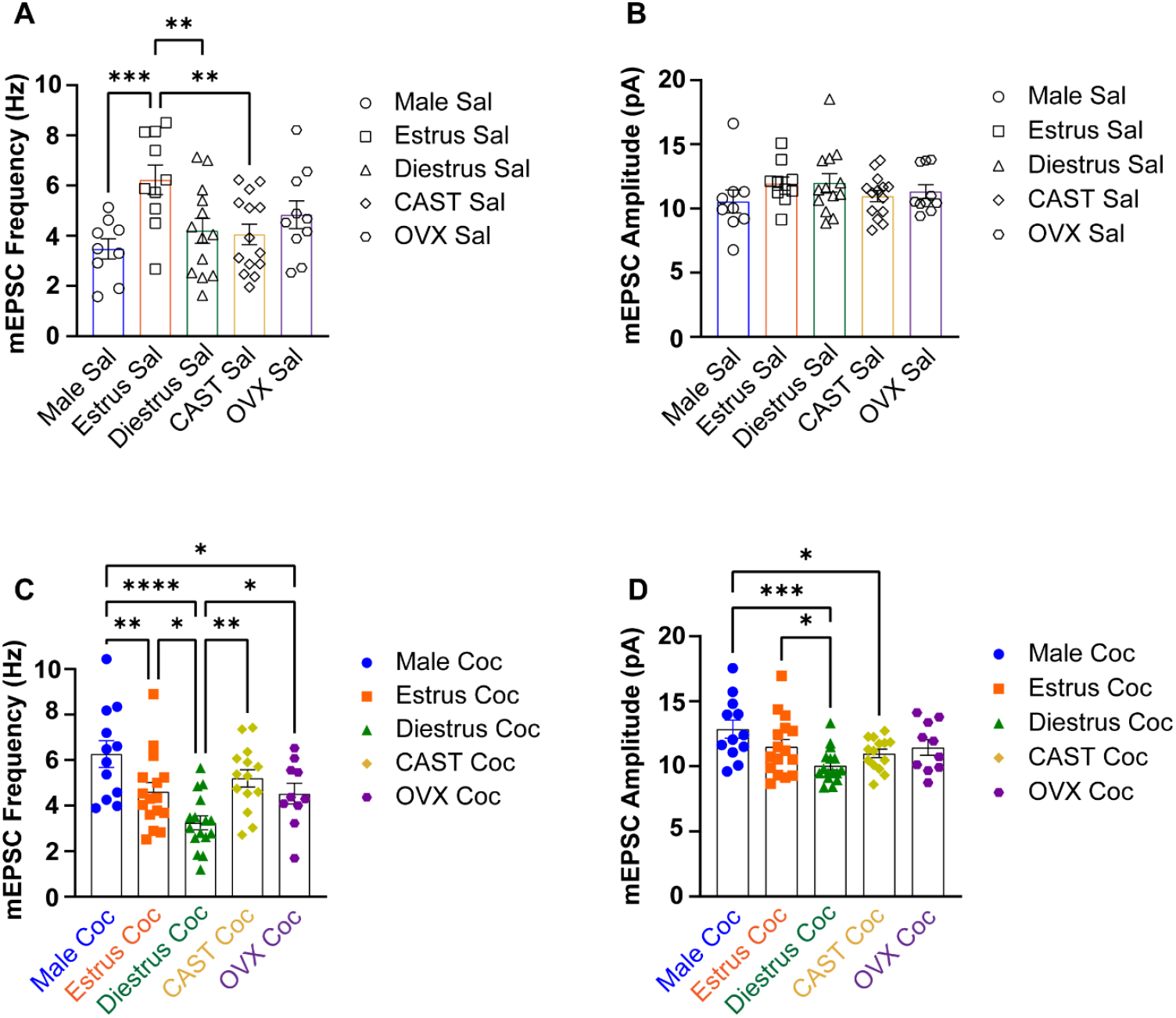
Gonadal hormones differentially impact mEPSCs. **A)** Summary NAcSh MSN mEPSC frequencies for saline-treated animals recorded during intact male (circle), estrus (square), diestrus (triangle), castrated (diamonds), and ovariectomized (hexagon). *p<0.05, **p<0.01, ***p<0.001. **B)** Summary NAcSh MSN mEPSC amplitudes for saline-treated animals recorded during intact male (circle), estrus (square), diestrus (triangle), castrated (diamonds), and ovariectomized (hexagon). **C)** Summary NAcSh MSN mEPSC frequencies for cocaine-treated animals recorded during intact male (blue, circle), estrus (orange, square), diestrus (green, triangle), castrated (mustard, diamonds), and ovariectomized (purple, hexagon). *p<0.05, **p<0.01, ***p<0.001, ****p<0.0001. **D)** Summary NAcSh MSN mEPSC amplitudes for cocaine-treated animals recorded during intact male (blue, circle), estrus (orange, square), diestrus (green, triangle), castrated (mustard, diamonds), and ovariectomized (purple, hexagon). *p,0.05, ***p<0.001. All neuronal recordings were obtained from n=3-6 mice. Abbreviations: saline (Sal), cocaine (Coc), Castrated (CAST), ovariectomized (OVX).

**Fig S6.**
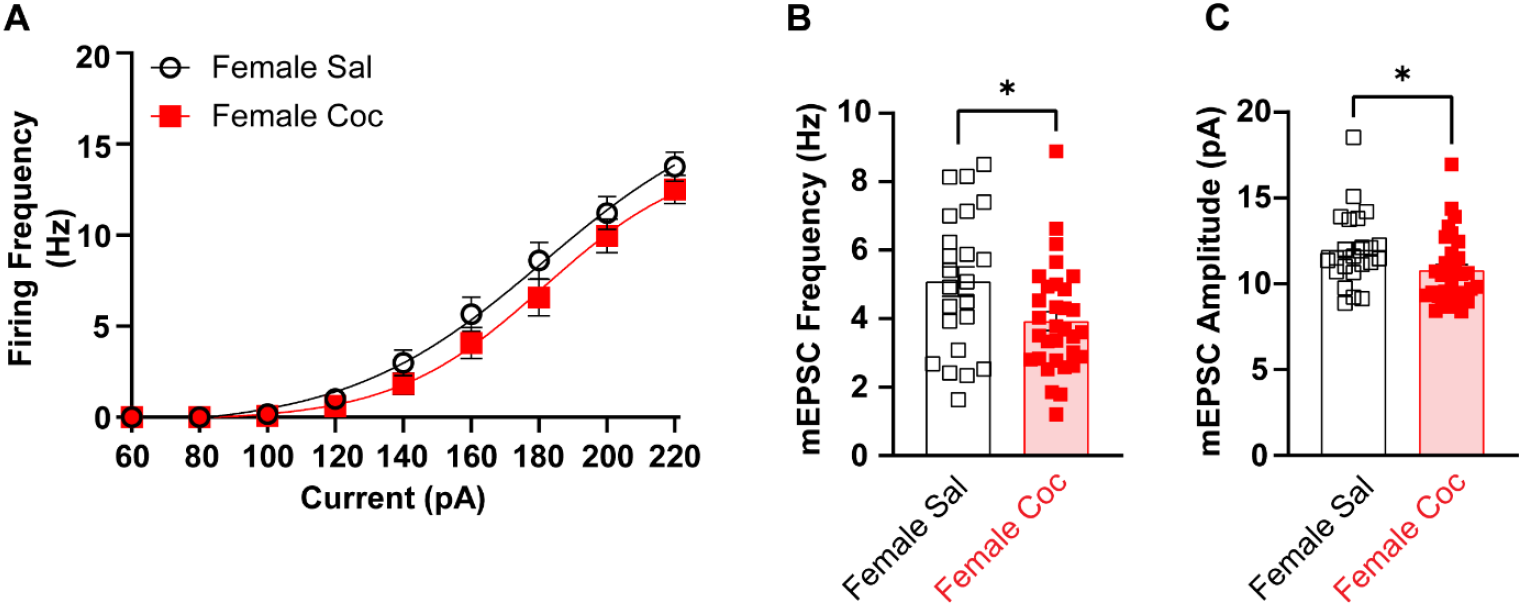
Neurophysiology of NAcSh MSNs when female data is pooled. **A)** Summary data showing the current injection response for combined female sal (black, open circles) vs combined female coc (red, closed circles). **B)** Summary data for mEPSC frequency from combined female sal (black, open squares) vs combined female coc (red, closed squares), *p<0.05. **C)** Summary data for mEPSC amplitude from combined female sal (black, open squares) vs combined female coc (red, closed squares), *p<0.05. Abbreviations: Saline (sal), cocaine (coc).

